# Compositional editing of extracellular matrices by CRISPR/Cas9 engineering of human mesenchymal stem cell lines

**DOI:** 10.1101/2023.07.28.550935

**Authors:** Sujeethkumar Prithiviraj, Alejandro Garcia Garcia, Karin Linderfalk, Bai Yiguang, Sonia Ferveur, Ludvig Nilsén Falck, Agatheeswaran Subramaniam, Sofie Mohlin, David Hidalgo, Steven J Dupard, Dimitra Zacharaki, Deepak Bushan Raina, Paul E Bourgine

## Abstract

Tissue engineering strategies predominantly rely on the production of living substitutes, whereby implanted cells actively participate in the regenerative process. Beyond cost and delayed graft availability, the patient-specific performance of engineered tissues poses serious concerns on their clinical translation ability. A more exciting paradigm consists in exploiting cell-laid, engineered extracellular matrices (eECM), which can be used as off-the-shelf materials. Here, the regenerative capacity solely relies on the preservation of the eECM structure and embedded signals to instruct an endogenous repair. We recently described the possibility to exploit custom human stem cell lines for eECM manufacturing. In addition to the conferred standardization, the availability of such cell lines opened avenues for the design of tailored eECMs by applying dedicated genetic tools. In this study, we demonstrated the exploitation of CRISPR/Cas9 as a high precision system for editing the composition and function of eECMs. Human mesenchymal stromal/stem cell (hMSC) lines were modified to knockout VEGF and RUNX2 and assessed for their capacity to generate osteoinductive cartilage matrices. We report the successful editing of hMSCs, subsequently leading to targeted VEGF and RUNX2-knockout cartilage eECMs. Despite the absence of VEGF, eECMs retained full capacity to instruct ectopic endochondral ossification. Conversely, RUNX2-edited eECMs exhibited impaired hypertrophy, reduced ectopic ossification and superior cartilage repair in a rat osteochondral defect. In summary, our approach can be harnessed to identify the necessary eECM factors driving endogenous repair. Our work paves the road towards the compositional eECMs editing and their exploitation in broad regenerative contexts.

## Introduction

Extracellular matrices (ECMs) are complex networks of proteins providing tissue structural and mechanical support, but also acting as growth factor storing and presenting entities^1^. As such, ECMs are receiving increasing attention in tissue engineering as templates capable of guiding homeostasis, remodeling, and regenerative processes^2,3,4^.

ECMs can be derived from native tissue or organs (native ECMs, nECMs) by applying a decellularization step which effectively removes the cellular fraction. The resulting nECMs are commonly used as biomaterials in regenerative medicine^5,6^, offering a natural biocompatible structure. However, they suffer from substantial batch-to-batch variation with their properties being largely affected by the tissue source and the decellularization process^5,7^. Importantly, while displaying a high level of biological complexity and fidelity, the composition of nECMs cannot be tailored to specific needs.

A valuable option emerged from recent advances in bioengineering which led to the development of synthetic ECMs (sECMs). These tunable materials are predominantly composed of biopolymers such as poly (lactic acid) (PLA), poly (glycolic acid) (PGA), poly (lactic co-glycolic acid) (PLGA) and Poly (ethylene glycol) (PEG), which can be functionalized with bioactive substances such as peptides, growth factors or enzymes^8,9,10^,. The possible modulation of sECM’s composition and chemistry offers precise control over their mechanical properties, degradability, and temporal release of instructive molecules towards improved tissue repair. Despite holding great promises, yet their structure and function remain simplified compared to their native counterpart ^11,12^.

With the aim of combining high biological fidelity and design flexibility, engineered ECMs (eECMs) have become a credible alternative. Those result from the exploitation of stem/progenitor populations capable of depositing ECMs under specific in vitro culture conditions^13,14,15,16^, which can be subsequently isolated from the cellular fraction and exploited as a cell-laid product. Early studies were conducted using eECM derived from primary cells, exhibiting variable performance associated with the inter-donor variability^17,18^.

Most recently, we demonstrated the possibility to standardize the production of eECMs through the engineering of dedicated human mesenchymal stromal/stem cell (hMSC) lines^19^. Precisely, the Mesenchymal Sword of Damocles – BMP2 (MSOD-B) line offered the generation of eECMs in the form of human cartilage templates^20^. Those cartilage templates can be further devitalized, lyophilized, and stored as an *off-the-shelf* tissue, retaining remarkable bone formation capacity by instructing endochondral ossification^21^.

The exploitation of cell lines as an unlimited cell source for eECM generation offers unprecedented standardization in graft production and performance. Meanwhile, it also consists of a robust cellular platform facilitating their further genetic engineering, to achieve the production of eECMs tailored in composition and function. Towards this objective, CRISPR/Cas9 appears as a versatile and efficient gene-editing tool. While this technology has been extensively used for genetic screening or disease modeling^22,23,24^, the possibility to edit hMSCs and study the compositional impact on deposited eECMs has not been investigated so far.

In this study, we aim at demonstrating that CRISPR/Cas9 can be applied as an editing tool for the engineering of custom eECMs. As proof-of-principle, our strategy relies on the CRISPR/Cas9-guided editing of the MSOD-B line for knocking out Vascular endothelial growth factor (VEGF) and Runt related transcription factor 2 (RUNX2), respectively a key pro-angiogenic protein and transcription factor critically involved in the endochondral ossification pathway. The resulting hMSC lines will be evaluated for their capacity to generate hypertrophic cartilage in vitro and instruct tailored skeletal tissue formation in vivo as cell-free and lyophilized templates. If successful, our study is expected to validate the use of CRISPR/Cas9 for eECMs edited in composition, while providing a novel platform towards decoding the necessary molecular signals driving effective tissue repair.

## Results

### CRISPR/Cas9 editing of hMSC lines lead to efficient VEGF knockout in cartilage tissues

VEGF is a known master regulator of angiogenesis^25^ and a key mediator of endochondral ossification. When reaching hypertrophy, chondrocytes highly express VEGF, prompting vasculature invasion and subsequent osteoprogenitor recruitment. This was proven to be essential for cartilage template remodeling into bone and bone marrow^26^. However, whether VEGF is a requirement to instruct ectopic endochondral ossification remains to be investigated. Here, we thus first aimed at engineering a hMSC line CRISPR/Cas9-edited for VEGF knockout and evaluate the corresponding impact on cartilage and endochondral bone formation (Figure 1A). To this end, we exploited the mesenchymal sword of Damocles bone morphogenetic type-2 (MSOD-B) as human cell line previously demonstrated as capable of endochondral ossification^20^ (Supplementary Figure 1A). Specific guide RNAs (gRNAs) targeting different regions of the VEGF gene but conserved across all isoforms of VEGF were designed based on a previously established protocol^27^ towards modifying the MSOD-B line. Three gRNAs targeting exon 1 (VEGF_1.1, VEGF_1,2, VEGF1.3), one targeting exon 2 (VEGF_2.1), and one targeting exon 8 (VEGF_8.1) were designed (Figure 1B, Supplementary Figure 1B). These gRNAs were cloned into the pU6-(BbsI)_CBh-Cas9-T2A-mCherry vector encoding the Streptococcus pyogenes Cas9 (SpCas9) machinery. Following transfection in MSOD-B cells, single cell clones were sorted based on the transient mCherry expression and expanded for characterization (Figure 1A). Out of 163 single clones, 14 could be successfully expanded (8.5 %). From these clones, five were retrieved from the gRNA 1.1, six from gRNA 1.3 and three from gRNA 8.1. In order to assess a successful editing and directly correlate it to a knockout of VEGF, an ELISA assay was performed to measure the concentration of VEGF in the supernatant of expanded MSOD-B clones (Figure 1C). In non-edited cells (MSOD-B, control), 5000 pg/mL were secreted and detected by ELISA. From the successfully expanded clones, most of them showed reduced but non-abolished secretion of VEGF. However, two clones exhibited undetectable levels of VEGF, both resulting from the gRNA targeting the Exon1 of the VEGF gene. These clones were defined as MSOD-BΔV1 and MSOD-BΔV2 and further characterized for their capacity to form cartilage.

**Figure 1.**
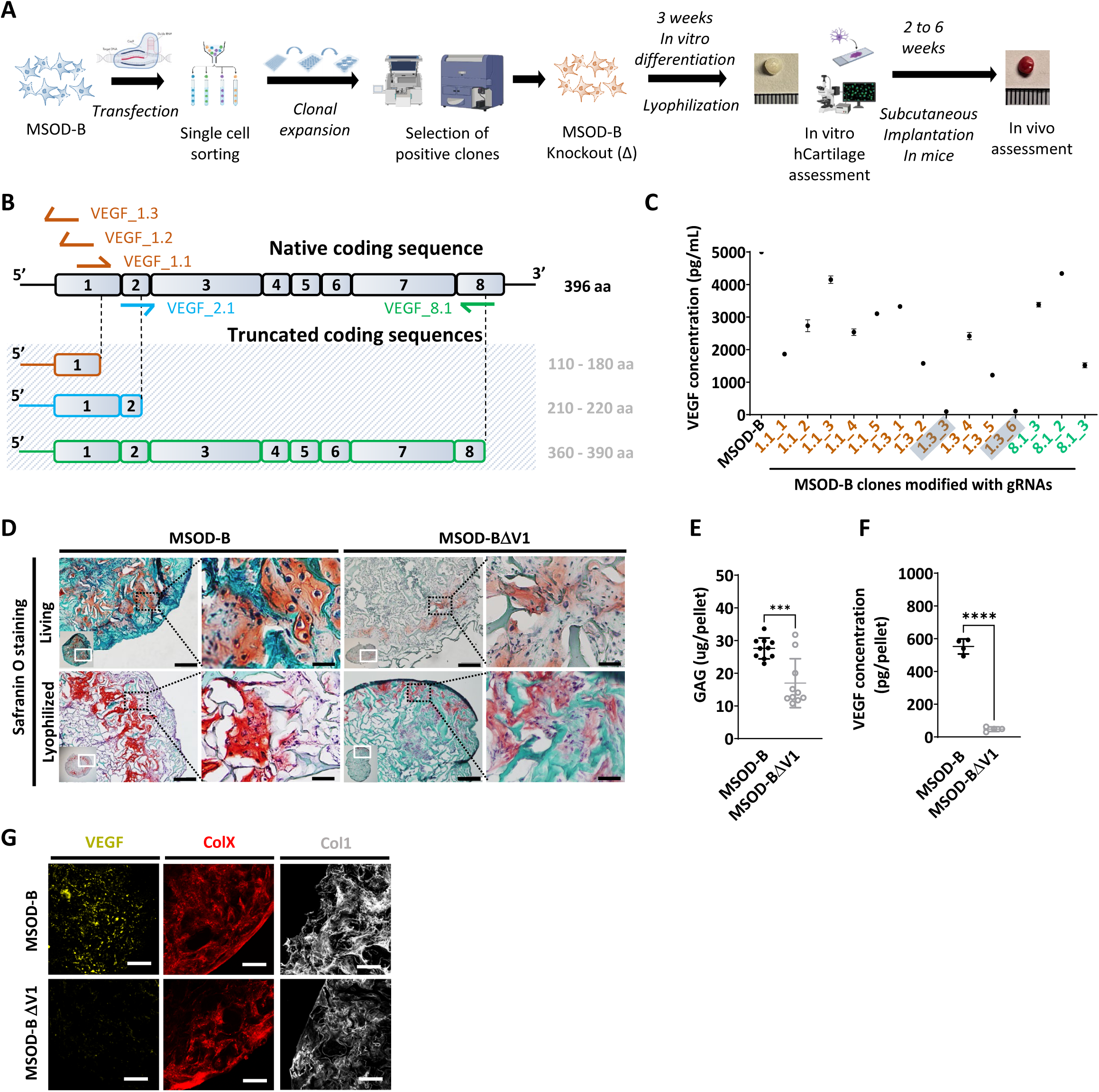
CRISPR/Cas9 editing of mesenchymal cell lines lead to efficient VEGF knockout in cartilage tissues. A) Experimental scheme depicting the generation of CRISPR/Cas9-edited MSOD-B lines, and the subsequent in vitro and in vivo tissue formation assessment. B) Overview of the native human VEGF coding sequence composed of eight exons. Designed gRNAs and their targeted binding sites are illustrated, as well as the corresponding expected impact on the coding sequence. gRNAs targeting exon 1 (orange) disrupt translation initiation and inhibit protein expression, gRNA targeting exon 2 (blue) disrupts VEGF receptor binding, and gRNA targeting exon 8 (green) alters the C-Terminal sequence and repress activation of protein. C) ELISA-based quantitative analysis of VEGF protein content in cell culture supernatant from expanded single cell colonies. From all clones, only two had no detectable level of VEGF (1.3_3 and 1.3_6). These clones were subsequently defined as MSOD-BΔV1 and MSOD-BΔV2. D) Histological assessment of living and lyophilized in vitro differentiated constructs using Safranin O staining (Scale bars = 100 µm and 20 µm for magnified areas). Both the MSOD-B and MSOD-BΔV1 displayed glycosaminoglycans (GAG) (orange to reddish in Safranin O), indicating successful cartilage formation (Scale bars = 200 µm). Left bottom inserts show the whole tissue section. E) Quantitative assessment of the total GAG content in MSOD-B and MSOD-BΔV1 in vitro differentiated constructs, post-lyophilization. Unpaired t-test, n = 10 biological replicates, ***p < 0.001. F) ELISA-based quantitative assessment of VEGF protein in in vitro differentiated constructs, post-lyophilization. Unpaired t-test, n = 3-4 biological replicates, ****p < 0.0001. G) Immunofluorescence images of MSOD-B and MSOD-BΔV1 tissues, post-lyophilization. Displayed images consist of 3D-stacks from 80-100 µm thick sections, stained for VEGF (yellow), Collagen Type X (COLX, red) and Collagen Type I (COL1, grey). A clear reduction in the VEGF signal could be observed in the MSOD-BΔV1 tissues, indicating a successful VEGF knockdown (Scale bars = 80 µm).

To this end, clones were expanded and seeded on collagen scaffold and induced for three weeks towards chondrogenic differentiation followed by lyophilization of the tissues. As anticipated with the MSOD-B tissues, histological assessment revealed the formation of a collagenous matrix with cartilage features (Safranin-O, Figure 1D). Similarly, a successful chondrogenic differentiation and deposition of glycosaminoglycans could be observed in the MSOD-BΔV1 (Figure 1D) constructs. A quantitative assessment (blyscan assay) confirmed the content in glycosaminoglycans in MSOD-B and VEGF-edited clones falling in the same concentration range although a lower amount was detected in the latest group (Figure 1E). Importantly, VEGF quantification in corresponding pellets validated the successful knockout of the protein, barely detectable in the MSOD-BΔV1 and ΔV2 tissues (47.64 pg/pellet in edited clones versus 552.7 pg/pellet in MSOD-B, Figure 1F).

To further examine the potential impact of VEGF-editing on tissue formation, we performed immunostaining analysis of the engineered cartilages. Confocal microscopy revealed strong deposition of Collagen Type I (COL1) and Collagen Type X (COLX) in all samples, characteristic of mature hypertrophic cartilage tissues (Figure 1G). However, while the immunostaining revealed the VEGF deposition in the MSOD-B ECM, the protein could not be detected in the MSOD-BΔV1 samples (Figure 1G).

Taken together, this data indicates the successful generation of MSOD-B lines knocked-out in VEGF. The MSOD-BΔV1 clones exhibited cartilage formation capacity with minimal level of VEGF. This validates the use of CRISPR/Cas9 as a precision tool to edit the composition of cartilage tissue.

### VEGF knockout cartilage tissues retain bone remodeling capacity despite reduced early-stage vascularization

To evaluate the functional impact of VEGF knockout in engineered constructs, we assessed both the angiogenic and bone formation assays. Since the two clones exhibited similar in vitro tissue formation capacity, only the MSOD-BΔV1 was selected for further performance assessment.

We first performed the Chorioallantoic Membrane (CAM) assay as ex vivo assessment of angiogenic potential (Figure 2A). This assay provides insights into the biological activity and neovascularization potential of each construct, through quantitative evaluation of vascular density around engineered grafts After 4 days, we observed an extensive vessel formation in the periphery of both MSOD-B and MSOD-BΔV1 lyophilized eECMs (Figure 2B). Quantification of vascular densities ^30^ suggested a reduced vessel formation in MSOD-BΔV1 samples but without reaching significance (Figure 2C and Supplementary Figure 2A). Thus, the CAM assay indicated that both MSOD-B and MSOD-BΔV1 cartilages retain angiogenic potentials.

**Figure 2.**
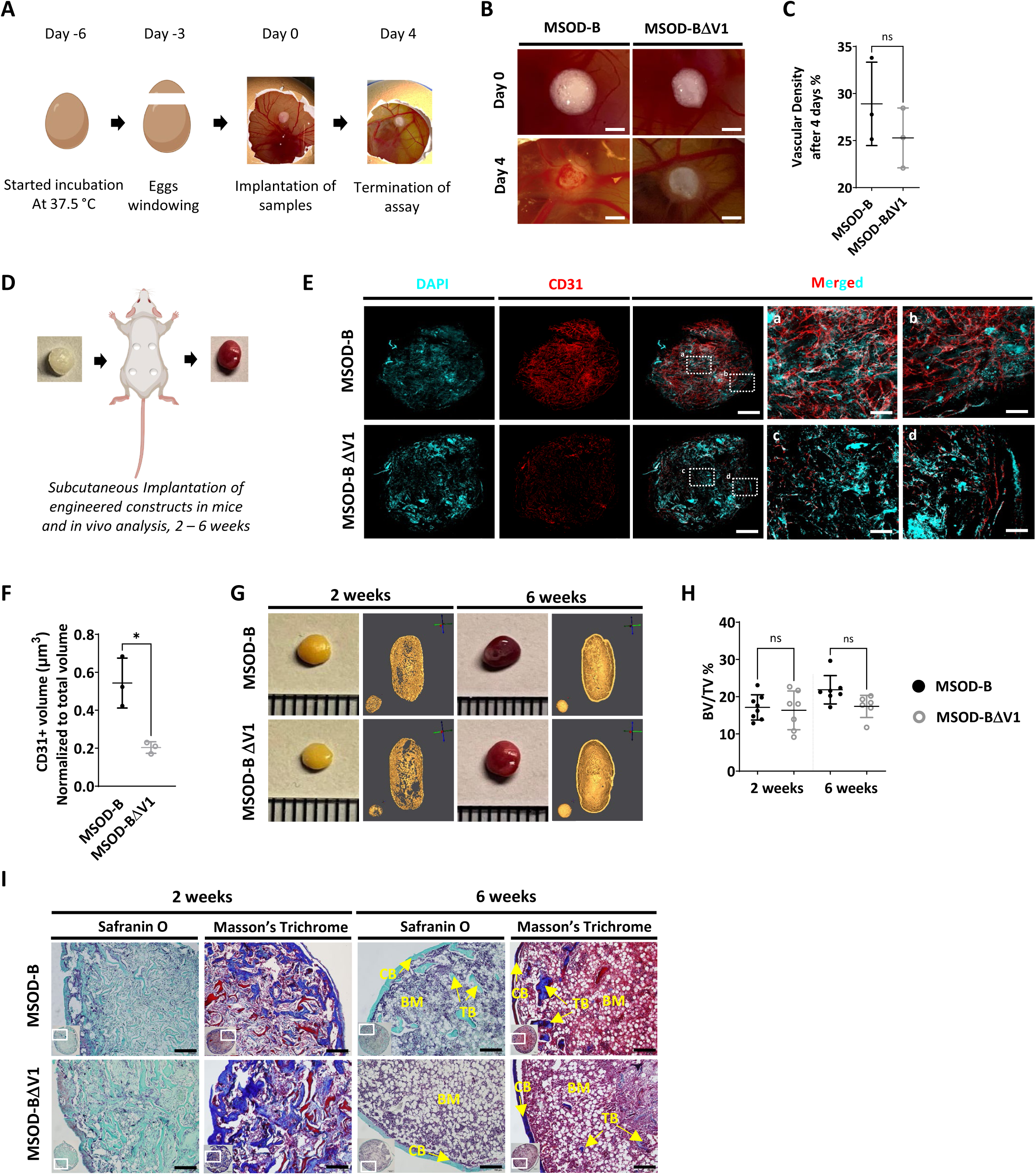
VEGF knockout cartilage tissues retain bone remodeling capacity despite reduced early-stage vascularization. A) Experimental scheme of the Chorioallantoic Membrane (CAM) assay, for ex vivo evaluation of angiogenic potential. B) Macroscopic comparison of in vitro MSOD-B and MSOD-BΔV1 constructs at day 0 (day of implantation) and day 4 (4 days post-implantation), illustrating robust de novo vessel formation perfusing the tissues (Scale bars at 1 mm). C) Quantitative Analysis of Vascular Density in MSOD-B and MSOD-BΔV1. Vascular densities were quantified from macroscopic images obtained on day 4 using ImageJ. Unpaired t-test, n = 3-4 biological replicates, n.s. = not significant. D) overview of the subcutaneous implantation procedure in mice and subsequent in vivo evaluation of vascularization from 2 to 6 weeks post-implantation. E) Immunofluorescence images of MSOD-B and MSOD-BΔV1 tissues two weeks post-in vivo implantation. Displayed images consist of 3D-stacks from 80-100 µm thick sections, vessels stained with mouse CD31 (red) and nuclei with DAPI (cyan) (Scale bars at 500 µm except for magnified white inserts at 80 µm). Box “a” and “c” display the periphery whereas Box “b” and “d” show the central region of MSOD-B and MSOD-BΔV1 constructs respectively. A reduction in tissue vascularization is observed in MSOD-BΔV1 samples. F) Quantitative analysis of the CD31 signal using an isosurface-based strategy (IMARIS software). Unpaired t-test, n = 3-4 biological replicates, *p < 0.05. G) Representative macroscopic and microtomography images of in vivo constructs retrieved at two and six weeks post-implantation. H) Microtomography-based quantification of the sample’s bone/ mineralized volume over their total volume (ratio). No significant differences between MSOD-B and MSOD-BΔV1 could be observed. (BV: Bone Volume, TV: Total Volume). Ordinary One-way ANOVA, n=8 biological replicates, n.s.=not significant I) Histological analysis of in vivo tissues using Safranin O and Masson’s trichrome stains, at two (2W) and six weeks (6W) post-implantation. Both sample types underwent full remodeling into a bone organ after 6 weeks, with presence of bone structures and a bone marrow compartment (Scale bars = 200 µm). CB – cortical bone; BM – bone marrow. TB – trabecular bone.

We next investigated in vivo the impact of the VEGF knockout on the graft capacity to undergo endochondral ossification, by implanting cartilage grafts subcutaneously in an immunodeficient mouse model. Importantly, prior to implantation, tissues were lyophilized using a pre-established protocol^31^. This allowed us to assess the performance of the generated tissue itself, in line with the idea of developing off-the-shelf substitutes. Samples were extracted after two and six weeks as early and late development timepoints (Figure 2D). After two weeks in vivo, we first evaluated the angiogenic potential of tissues by quantitative confocal microscopy imaging of 100 µm thick sections stained for CD31, a well-defined vascular marker. This allowed us to evidence a dense vascular network in MSOD-B constructs, covering the entirety of the grafts (Figure 2E). Instead, MSOD-BΔV1 displayed a more limited vascularization, predominantly at the tissue periphery (Figure 2E). Quantification confirmed these observations with MSOD-B exhibiting a significantly higher volume of vessels per total section volume (0.523 µm^3^ versus 0.231 µm^3^ for MSOD-B and MSOD-BΔV1 respectively, Figure 2F).

Using microtomography, we further assessed the amount of mineralized tissue formed in a temporal fashion. At two weeks, the formation of a cortical ring could already be observed in both MSOD-B and MSOD-BΔV1 tissues (Figure 2G). Quantifications revealed a similar bone/mineralized volume normalized to the total volume of the graft (BV/TV, Figure 2H, supplementary figure 2B and 2C) with 17 % and 16 % in MSOD-B and MSOD-BΔV1 samples respectively. At the six-week timepoint, trabecular structures could be observed in reconstructed 3D scans from both groups (Figure 2G). Remarkably, histological analysis confirmed the similar development of the tissues with bone formation (Massońs trichrome) already at two weeks post-implantation and minimal remnants of cartilage (Safranin O) (Figure 2I). At six weeks both samples remodeled into fully mature bone organs, characterized by presence of cortical and trabecular structures as well as a bone marrow tissue filling the cavity (Figure 2I).

These results indicate that the absence of VEGF in cartilage tissue can delay the early vascularization of MSOD-BΔV samples. However, this did not prevent nor impact the remodeling of the lyophilized grafts into bone and bone marrow tissues indicating that VEGF is non-essential in order to efficiently instruct endochondral ossification.

### RUNX2 knockdown does not prevent chondrogenic differentiation but impairs hypertrophy

We next investigated whether the composition and thus function of a graft could be modified by editing transcriptional factors involved in mesenchymal cell differentiation. Using CRISPR/Cas9, we thus targeted the RUNX2, a known master regulator particularly important for chondrocyte differentiation and hypertrophy^32,33^. We designed gRNAs targeting the coding regions involved in cell signaling integration, activation, and inhibition domain which are conserved in both isoforms of RUNX2 (Figure 3A, Supplementary Figure 3A).

**Figure 3.**
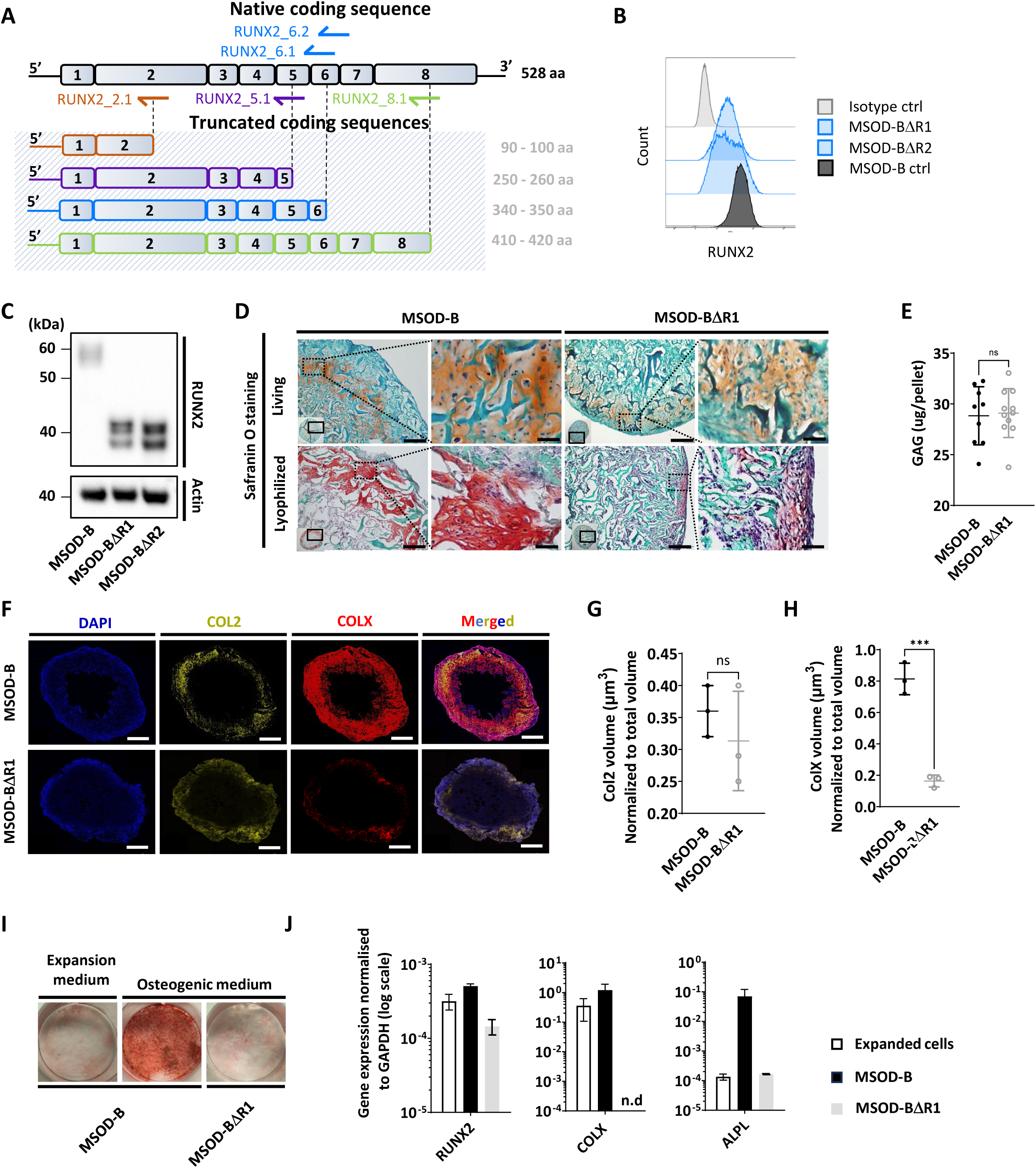
RUNX2 knockout does not prevent chondrogenic differentiation but impairs hypertrophy. A) Overview of the human RUNX2 coding sequence comprising 8 exons. gRNAs and their corresponding expected protein structure. The gRNA targeting exon 2 (orange) disrupts the DNA binding domain, gRNA targeting exon 5 (violet) disrupts nuclear translocation, gRNAs targeting exon 6 (blue) disrupt the transcriptional activation domain and gRNAs targeting exon 8 (green) disrupt the nuclear matrix targeting signal and repress the protein function. B) Intracellular flow cytometry for RUNX2 detection in MSOD-B and RUNX2-edited clones. A clear protein reduction could be observed in the 6.1_1 and 6.1_23 clones. C) Western blot analysis of RUNX2 in cultured MSOD-B and RUNX2-edited cells. The genetic editing of RUNX2 is confirmed by the detection of the truncated proteins. Actin is used as a control to normalize the protein content. D) Histological analysis of living and lyophilized in vitro differentiated constructs using Safranin O staining (Scale bars = 100 µm and 20 µm for magnified areas), indicating the presence of cartilage matrices. E) Quantitative assessment of the total GAG content in corresponding in vitro generated lyophilized tissues. Unpaired t-test, n=10-11 biological replicates **p < 0.01. F) Immunofluorescence images of MSOD-B and MSOD-BΔR1 lyophilized tissues. Displayed images consist of 3D-stacks from 80-100 µm thick sections, stained for DAPI (blue), Collagen Type II (COL2, yellow) and Collagen Type X (COLX, red). A clear reduction in the COLX signal could be observed in the MSOD-BΔR1 tissues, indicating impaired hypertrophy. (Scale bars = 500 µm) G) Isosurface-based quantification of the COL2 immunofluorescent signal using the IMARIS software. No significant difference between groups confirm the retention of cartilage formation in RUNX2-edited constructs. Unpaired t-test, n=3 biological replicates, n.s.=not significant. H) Isosurface-based quantification of the COLX immunofluorescent signal using the IMARIS software, confirming the disruption of hypertrophy in the MSOD-BΔR1 constructs. Unpaired t-test, n=3, ***p < 0.001. I) Alizarin Red staining evidencing a lack of mineralization in the MSOD-BΔR1 culture compared to the MSOD-B. J) Quantitative PCR analysis displaying the relative expression levels of osteogenesis-related genes: RUNX2, COL1, and ALPL. The expression is normalized to GAPDH as housekeeping gene. n.d= not detected.

One gRNA was designed for targeting exon 2 (RUNX2_2.1), one targeting exon 5 (RUNX2_5.1), two targeting exon 6 (RUNX2_6.1 and RUNX2_6.2) and one targeting exon 8 (RUNX2_8.1). Similar to the VEGF set-up, gRNAs were transfected in MSOD-B cells together with a pU6-(BbsI)_CBh-Cas9-T2A-mCherry plasmid to ensure transfection efficiency and single cell sorting of positive clones. Out of 385 single clones, 62 could be successfully expanded, corresponding to a 16.1% efficiency. From these clones, five were derived from the RUNX2_5.1 gRNA, 18 from RUNX2_6.1, 15 from RUNX26.2 and 24 from RUNX2_8.1.

To identify successfully edited clones, we first screened for those exhibiting a decrease in their RUNX2 protein expression using intra cellular flow cytometry. Among the 62 clones, 17 displayed a reduced RUNX2 pattern as compared to the MSOD-B control (Figure 3B and Supplementary Figure 3B) and were further sent for sequencing analysis. This led to the identification of two successfully edited clones, demonstrating a point mutation in the exon 6 of RUNX2 gene (Supplementary Figure 3C). To confirm the knockout impact on the transcription factor structure, a Western Blot analysis was conducted on cellular extracts of in vitro cultured cells. MSOD-B cells exhibited intact RUNX2 proteins of 52 – 62 kDa^34,35^(Figure 3C). Instead, the RUNX2-edited cells displayed truncated versions of 32 – 42 kDa, consistent with the expected mRNA shortening.

These two clones were defined as MSOD-BΔR1 and MSOD-BΔR2 and further assessed for their ability to form cartilage in vitro. Following 3D chondrogenic differentiation, samples were lyophilized and processed for histological analysis. The presence of cartilage structures embedded in a collagenous matrix could be observed in all MSOD-B, MSOD-BΔR2 (Figure 3D) and MSOD-BΔR1 tissues (Supplementary Figure 3D). This was confirmed quantitatively using the Blyscan assay, with detectable glycosaminoglycans in all groups (Figure 3E, Supplementary Figure 3E). Using immunostaining combined with quantitative confocal microscopy, the possible impact of RUNX2-editing on tissue hypertrophy was further investigated. First, Collagen Type II (COL2) staining confirmed the distribution of cartilage matrix across MSOD-B, MSOD-BΔR1 and MSOD-BΔR2 samples (Figure 3F and Supplementary Figure 3F). However, we observed a clear distinction in the Collagen Type X (COLX) expression pattern, a specific marker of hypertrophy which was hardly detectable in MSOD-BΔR tissues (Figure 3F and Supplementary Figure 3F). Subsequent quantification confirmed a significant reduction of COLX volume in MSOD-BΔR1 (0.236 µm^3^) and MSOD-BΔR2 (0.182 µm^3^) compared to MSOD-B (0.81 µm^3^) (Figure 3G, Figure 3H, Supplementary figure 3G-H-I).

To further characterize the functional impact of the RUNX2 knock-down, the MSOD-B and MSOD-BΔR1 osteogenic differentiation capacity was assessed in vitro. After 3 weeks of culture in osteogenic medium (or expansion medium, control), Alizarin Red staining revealed a s a marked absence of mineralization in MSOD-BΔR1 culture, while in stark contrast with MSOD-B cells (Figure 3I). Quantitative PCR analysis indicated a lower expression of COLX and ALPL in MSOD-BΔR1 (Figure 3J), key osteogenic-associated genes, of interest, the expression level of RUNX2 was only partially decreased in MSOD-BΔR1, in line with the truncated mRNA reducing but not abrogating qPCR primers binding probability.

In summary, we here report the successful RUNX2 knockout in MSOD-B lines using CRISPR/Cas9. RUNX2 did not impair chondrogenesis but prevented the tissue hypertrophy, as well as the osteogenic potential of the cells.

### RUNX2 knockout in cartilage tissues disrupts effective ectopic bone formation

To assess the corresponding impact of RUNX2-edited cartilages on bone formation, MSOD-B tissues (as control) and MSOD-BΔR1 were lyophilized and implanted subcutaneously in immunodeficient mice for two to six weeks. From microtomography image reconstructions, we observed an evident reduction of mineralization in MSOD-BΔR1 and compared to MSOD-B (Figure 4A) already after two weeks. This qualitative and visual difference in mineralization persisted after six weeks in vivo (Figure 4A). Subsequent quantifications confirmed these observations with a clear reduction in BV/TV as early as week 2 (16.46 % in MSOD-B and 7.01 % in MSOD-BΔR1) and persisting at week 6 (20.03% in MSOD-B and 4.25 % in MSOD-BΔR1) (Figure 4B, Supplementary Figure 4A, Supplementary Figure 4B). Histological analysis revealed an early bone formation at two weeks in both samples, but to a lower extent in the MSOD-BΔR1 group (Figure 4C). As anticipated, MSOD-B tissues underwent full remodeling after six weeks in vivo. In sharp contrast, the MSOD-BΔR1 only displayed a partial maturation with limited presence of bone and bone marrow.

**Figure 4.**
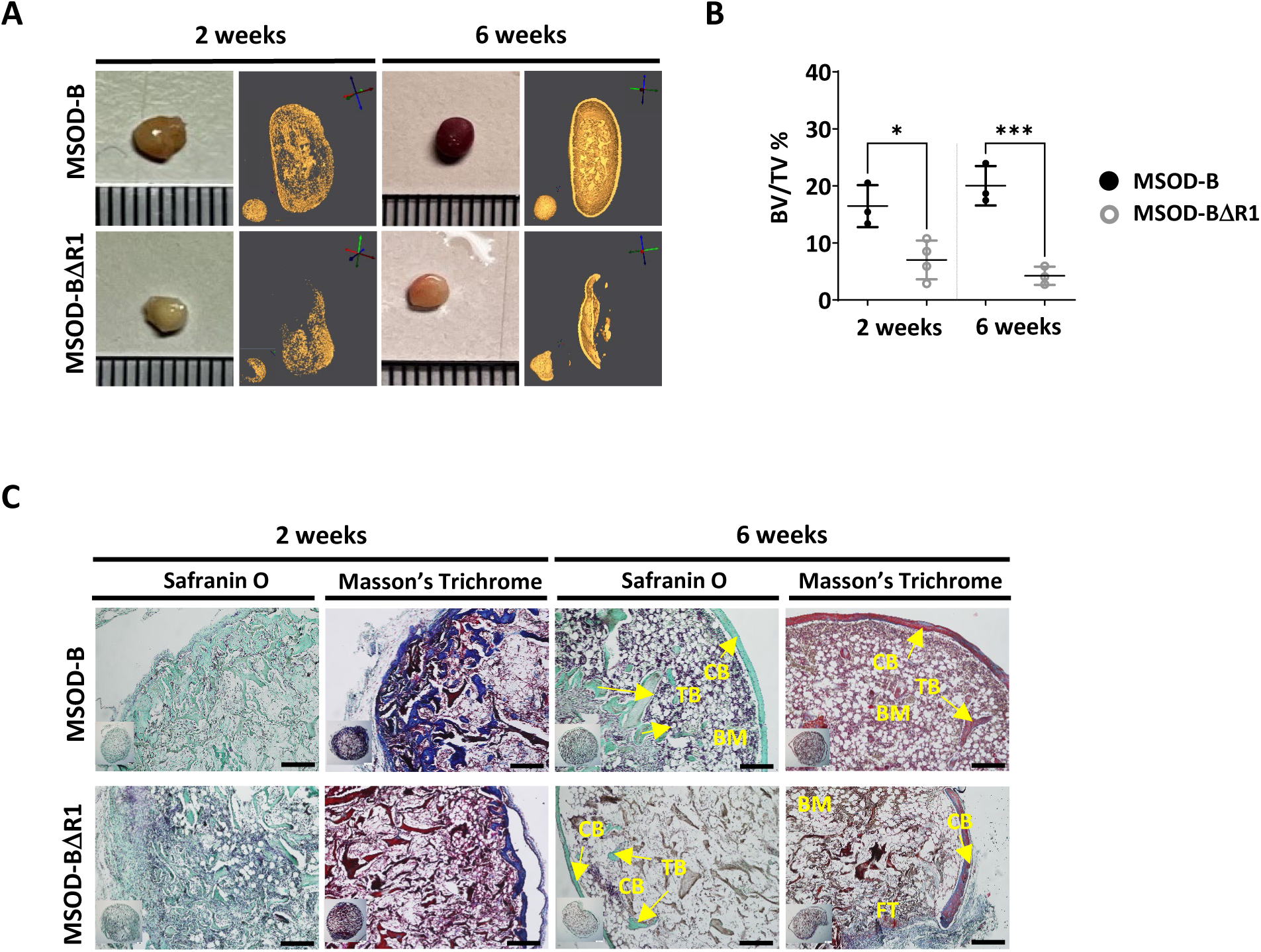
RUNX2 knockout in cartilage tissues disrupts effective ectopic bone formation. A) Representative macroscopic and microtomography images of in vivo constructs retrieved at two and six weeks post-implantation. B) Microtomography-based quantification of the sample’s bone/mineralized volume over their total volume (ratio). (BV: Bone Volume, TV: Total Volume). A marked difference is observed as early as two weeks, with a clear lower mineral content in MSOD-BΔR1 samples. Ordinary one way ANOVA, n=3 biological replicates, *p < 0.05, ***p < 0.001. C) Histological analysis of in vivo constructs using Safranin O and Masson’s trichrome stains. After two weeks (2W), a higher bone formation is already evident in the MSOD-B control group. The MSOD-BΔR1 samples explanted after 6 weeks (6W) displayed presence of cortical and trabecular bone, but also large amount of fibrous tissue indicating an incomplete remodeling. (Scale bars = 200 µm).

Altogether, this indicates an incomplete remodeling of RUNX2 edited samples with significantly delayed cortical and trabecular structure formation. This correlates with the impaired hypertrophic phenotype in the corresponding in vitro samples.

We next investigated a potential batch-to-batch variability in the generation of engineered cartilage tissues using the various MSOD-B lines. For each modified lines, independent batches were generated and the amount of glycosaminoglycans was assessed in resulting lyophilized tissues. This revealed a rather consistent generation of cartilage across batches, with no statistical differences across groups (Supplementary Figure 5A, Supplementary Figure 5B, Supplementary Figure 5C). We further conducted a quantitative assessment of pellet volume variability across tissues (Supplementary Figure 5D), showing minimal differences between the groups, indicating a limited size variation across the samples. Taken together, this points at a low variability across batches of cartilage grafts displaying comparable volume and GAG content.

### RUNX2 knockout in cartilage tissues leads to better cartilage regeneration in a rat osteochondral defect

In order to assess the performance of CRISPR-Cas9 edited eECMs in a relevant skeletal regenerative context, a proof-of-concept study in an immunocompetent rat osteochondral defect was performed. In addition to the lyophilization process, the MSOD-B and MSOD-BΔR1 tissues were also decellularized following a pre-established protocol^37^ in order to reduce inflammation resulting from the rat immune system. We performed a quantitative assessment of the total GAG content in decellularized MSOD-B and MSOD-BΔR1 constructs, showing partial preservation of GAG in the two groups compared to their living counterparts (Supplementary Figure 6A). We further assessed the total DNA content in MSOD-B and MSOD-BΔR1 constructs before and after decellularization as a measure of decellularization efficiency (supplementary figure 6B). Post-decellularization, the DNA content was significantly reduced to around 540 ng/pellet for Runx2-modified constructs and 280 ng/pellet for MSOD-B grafts (supplementary figure 6B), accounting for a reduction of 98.1% and 97.8% in DNA content respectively indicating an efficient decellularization process across samples. The tissues were subsequently placed in the sub-chondral defect in the rat distal femur. The defect consisted in a drill hole of 1mm diameter and 2mm depth (Figure 5A and supplementary figure 6C). Positive controls consisted of untreated healthy rats.

**Figure 5.**
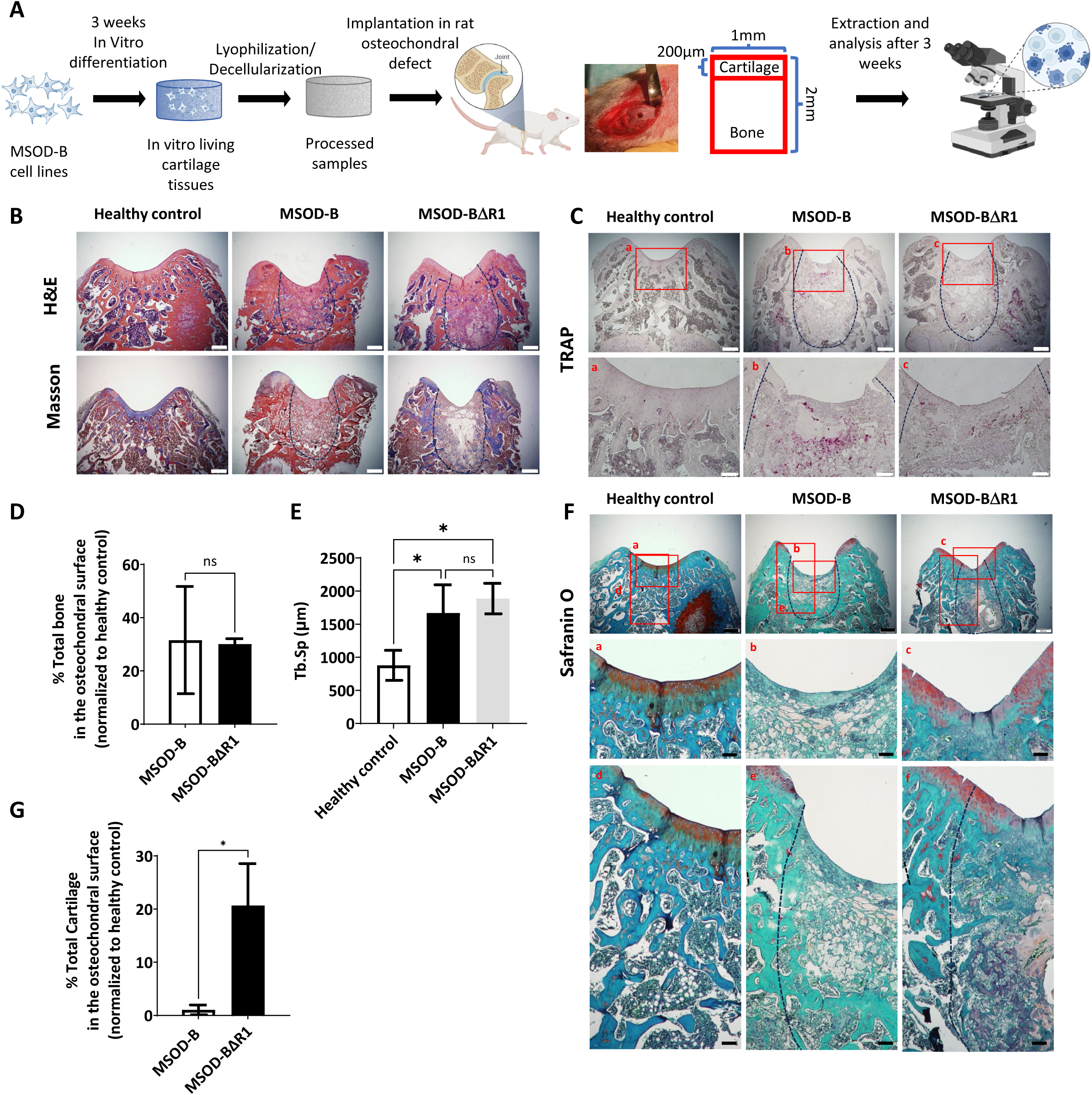
RUNX2 knockout in cartilage tissues leads to better cartilage regeneration and maintenance in osteochondral defect in rats. A) Experimental scheme for the regenerative potential assessment of MSOD-B & MSOD-BΔR1 cartilage tissues in a rat osteochondral defect. B) Histological analysis of the osteochondral defects for each group using H&E and Masson’s trichrome stains, at three weeks post-implantation. The dash-line marks the defect area. (Scale bars = 500 µm) C) Histological analysis the osteochondral defects using TRAP staining reporting osteoclastic activity, at three weeks post-implantation. The dash-line marks the defect area. (Scale bars = 500 µm and 100 µm for magnified areas) D) Microtomography-based quantification of the sample’s total bone/mineralized volume normalized to the healthy control in percentage. Unpaired t-test, n=3 biological replicates, n.s.=not significant. E) Image J based quantification of Trabecular separation (Tb.Sp). One way ANOVA test, n=3 biological replicates, *p < 0.05 F) Histological analysis of the osteochondral defects using Safranin O staining. After three weeks, a higher regeneration of the surface cartilage is evident in the MSOD-BΔR1 group (a,b,c) (Scale bars = 500 µm and 100 µm for magnified areas). The magnified regions of the subchondral area (d,e,f) show higher cartilage remnants and integration in the MSOD-BΔR1 group. G) Quantitative analysis of cartilage regeneration in the osteochondral surface as compare to the healthy control (100%). Unpaired t-test, n=3 biological replicates, *p < 0.05.

Following explantation, histological stainings and micro-tomography were performed on all sample groups. Hematoxylin and Eosin (H&E) and Masson’s Trichrome indicated de novo bone formation in the defect area for both the MSOD-B and MSOD-BΔR1 groups (Figure 5B and supplementary figure 6D). The presence of fibrotic tissue as well as a reduced marrow compartment suggested an incomplete remodeling at that timepoint (Figure 5B). Upon Tartrate-resistant acid phosphatase (TRAP) staining, we observed elevated osteoclastic activitiy in MSOD-B samples compared to MSOD-BΔR1 potentially indicating a reduced rate of active bone formation in MSOD-BΔR1 (Figure 5C). The micro-CT quantification revealed a 30% repair of the damaged bone in the defect for both MSOD-B and MSOD-BΔR1 groups (Figure 5D, Supplementary figure 6E). Of interest, a significantly higher trabecular separation was observed in MSOD-B and MSOD-BΔR1, confirming an ongoing bone remodeling process (Figure 5E). Using the Image J software, we quantified the trabecular thickness (tb.Th) in MSOD-B and MSOD-BΔR1 constructs. The analysis revealed no significant differences in tb.Th between the groups, indicating that trabecular architecture remained consistent across the samples (supplementary figure 6F).

While no statistical differences in bone formation could be identified between the two groups, the MSOD-BΔR1 tissue led to a detectable regeneration of the cartilage area, as revealed by Safranin-O staining (Figure 5F) in the chondral zone (Figure 5F, a,b and c magnification). Interestingly, the MSOD-BΔR1 group exhibited a higher remnants of GAGs in the subchondral area and better integration (Figure 5F, d,e,f magnification) to the host tissue. Quantification of cartilage tissue within condyle surface area confirmed a poor-to-no repair in the MSOD-B group (1.05%, Figure 5G). In sharp contrast, the MSOD-BΔR1 implanted eECMs initiated a chondral regeneration reaching approximately 20.67% of the total healthy cartilage area. The integration and regeneration potential of engineered constructs was further evaluated by performing a semi-quantitative histological assessment following a pre-existing grading system^39^ (Table 1 and Supplementary table 1). This approach allows for the systematic evaluation of critical repair tissue parameters, offering a comparative measure of the regenerative efficacy of the engineered constructs against healthy tissue benchmarks. In the assessment of stained femur condyle sections, cellular morphology and matrix staining of the MSOD-BΔR1 group (66% and 50%, respectively) were significantly better compared to the MSOD-B one (33% and 16%, respectively). Although both constructs exhibited reduced cartilage thickness and subchondral bone regeneration, MSOD-BΔR1 consistently outperformed the MSOD-B grafts (33.33% versus 8.33% and 25% versus 16.66%). Surface regularity and integration (Figure 5F) with adjacent cartilage also revealed better outcomes in MSOD-BΔR1 (41% and 50%) as opposed to MSOD-B (50% and 33%), further indicating an enhanced regenerative potential of the MSOD-BΔR1 constructs.

**Table 1:** Scoring rate for individual categories. Semi-quantitative analysis utilizing a modified histological grading system, including parameters such as cellular morphology, matrix staining, surface regularity, thickness of cartilage, subchondral bone formation and integration of adjacent cartilage, resulting in a comprehensive assessment of tissue regeneration and integration

| Scoring Rate (%) (SD) |  |  |  |  |  |  |
| --- | --- | --- | --- | --- | --- | --- |
|  | Cell Morphology | Matrix Staining | Surface Regularity | Thickness of Cartilage | Regenerated Subchondral Bone | Integration with Adjacent Cartilage |
| **Healthy** | 100 (0) | 100 (0) | 100 (0) | 100 (0) | 100 (0) | 100 (0) |
| **MSOD-B** | 33,33 (28,86) | 16,66 (14,43) | 50 (0) | 8,33 (14,33) | 16,66 (14,33) | 33,33 (14,33) |
| **MSOD-B ΔR1** | 66,66 (14,33) | 50 (0) | 41,66 (14,33) | 33,33 (28,86) | 25 (25) | 50 (0) |

In conclusion, while both grafts yielded comparable bone regeneration within the osteochondral defects, only the RUNX2-deleted grafts supported a cartilage regeneration.

## Discussion

We here report the possibility to edit the composition and function of eECM by CRISPR/Cas9 engineering of human mesenchymal lines. VEGF knockout led to successful in vitro formation of cartilage with targeted protein depletion. Upon implantation, this impacted the graft vascularization onset but not the endogenous instruction of the endochondral program. In turn, RUNX2 knockout prevented cartilage hypertrophy in vitro, significantly delaying ectopic bone and bone marrow formation in vivo. This strategy was validated in a functional osteochondral defect model, whereby prevention of cartilage hypertrophy by RUNX2 deletion improved cartilage regeneration.

A clear interdependency of angiogenesis and osteogenesis occurs upon bone formation^40,41^. For this reason, VEGF-enrichment has been naturally proposed as a strategy to accelerate or increase bone graft vascularization^42,43^. Our study demonstrates that VEGF knocked-out cartilage templates retained full osteoinductive potential in a challenging ectopic environment. Those results are in sharp contrast with intramembranous strategies^44,45^, including our previous work^19^, whereby grafts remain insufficient in promoting complete bone remodeling even upon VEGF-enrichment. This suggests that the first stage of host progenitor recruitment by our eECM is not VEGF dependent, although subsequent tissue vascularization and remodeling may be orchestrated in a second step by endogenous cell secretion. Other pro-angiogenic proteins embedded in the matrix may also have compensated for the lack of VEGF, such as BMP-2^20^ known to stimulate endothelial cell proliferation, migration, and differentiation^46,47^.

Conversely, RUNX2 deletion was shown to fully prevent cartilage hypertrophy, in line with published mouse models also reporting a lack of endochondral ossification and lethality at birth upon RUNX2 knockout^48,49^. Here, MSOD-BΔR cartilage templates still exhibited osteoinductive capacity, but with cortical and trabecular structures largely reduced as compared to controls. Future work may clarify if this strictly results from the absence of COLX, or if an additional specific cartilage matrix component (e.g. matrix metalloproteinases) impaired the instruction of endochondral ossification.

The performance in a regenerative context was further evaluated in an immunocompetent rat osteochondral defect model^50^. We defined an early timepoint of 3-week as providing the opportunity to assess the early contribution of the grafts to both the cartilage and bone tissue regeneration. Strikingly, only the MSOD-BΔR could contribute to neo-chondrogenesis. We hypothesize that the absence of hypertrophic features prevented the template degradation and favored its integration to native cartilage Despite the repair performance being limited to an approximate 20% of native cartilage restoration, this is remarkable in light of the BMP-2 content in our ECM prompting endochondral ossification ^51,52,53^. In fact, preventing the in vivo remodeling of engineered cartilage templates remains challenging in the skeletal regeneration field, with bone marrow MSCs (BM-MSCs) cartilage systematically reaching hypertrophy and subsequent ossification. The molecular mechanisms remain nonetheless elusive, warranting further studies comprising additional timepoints to decipher the repair dynamic as well as the long-term stability of newly formed tissue. It also prompts a comparison with other recently proposed *off-the-shelf* strategies, based on ECM^54^ or synthetic scaffold materials^55^, in order to comprehend the feasibility to exploit our grafts for stable cartilage / joint repair^56,57^. Taken together, while the relevance of CRISPR/Cas9 eECMs in cartilage repair remains to be further validated, our study demonstrates the relevance of genetically-edited ECMs in regenerative contexts.

CRISPR/Cas9 editing of human mesenchymal cells has previously been explored for gene and cell therapy applications^58,59^, for tailoring the immunomodulatory and/or differentiation capacity of engineered cells. However, no studies have described the exploitation of editing strategies for the generation of eECMs, whereby modified cells are absent from the final grafting product. Our work illustrates that this can be achieved by direct targeting of secreted factors typically embedded upon ECM deposition, as demonstrated with VEGF. Alternatively, we also propose the knockout of key transcription factors as a strategy for custom eECM generation by impacting their tissue developmental/maturation stages.

A clear implication of our work lies in the possibility to decipher the necessary factors capable of instructing de novo tissue formation. Those findings will be of high relevance for the design of eECMs tailored in composition. Beyond bone repair, our study also bears high relevance in other regenerative contexts. In fact, an exciting opportunity also lies in harnessing CRISPR/Cas9 for editing eECMs and tuned their immunogenicity. Key inflammatory components could be turned down, leading to improved efficacy of repair in line with the immuno-engineering principles^60,61^.

Importantly, our study describes the exploitation of eECMs in a lyophilized form, thus conferring an off-the-shelf storage solution. While the lyophilization and decellularization process can affect the ECM integrity, the resulting grafting products were demonstrated to retain regenerative properties. Together with the standardization of production conferred by stable cell sources, our concept offers exciting translational opportunities. In fact, instrumental to this work is the use of dedicated hMSC lines. The genetic modification of primary cells is laborious, and their limited lifespan ex vivo challenges their selection, characterization, and exploitation for tissue engineering applications^62,63^. Here, the MSOD-B was harnessed as an unlimited cell source with robust differentiation potential. This confers a higher standardization potential, although cell line-derived product can also be subject to batch dependency. Our study reports a limited batch-to-batch variation but a stringent characterization of cell lines capacity may be required upon substantial passage. The editing of eECMs remains tedious in part due to the limited CRISPR/Cas9 efficiency and potential off-target effects, calling for a systematic clonal selection. In addition, while large CRISPR/Cas9 screening can be performed in other stem cell systems^64,65^, the critical cell mass required for effective cartilage formation affects parallelization and leads to limited throughput. Nonetheless, after the identification and characterization process, the resulting cell lines can be banked and used for unlimited tissue manufacturing.

The MSOD-B line remains the only human cell source capable of priming endochondral ossification by engineering living or cell-free grafts. This was demonstrated to be driven by the combined low dose of BMP-2 embedded in the tissue (∼40 ng/tissue) together with glycosaminoglycans and other thousands of identified ECM proteins^20^. The BMP-2 amount is thus far below the typical amount used in sECMs approaches for effective osteoinduction, falling in the microgram to milligram range^31^. This is one key advantage of eECM graft, exhibiting a biological complexity that so far has not been matched by sECMs. Ideally, the two approaches are complementary as the editing of eECMs could inform on the necessary but sufficient factors driving effective regeneration. Those would in turn be embedded in a sECMs strategy and offer a fast and possibly cost-effective solution.

To conclude, the present work offers a platform for decoding factors involved in tissue regeneration and generate tailored eECM. Here illustrated in the context of skeletal repair, our study may offer similar opportunities in other regenerative situations.

## Materials and Methods

### Cell expansion

MSOD-B cells and their modified progeny were cultured in a humidified incubator at 37 °C and 5% CO_2_ using complete medium consisting of α-minimum essential medium (αMEM) supplemented with 10% fetal bovine serum, 1% HEPES, 1% sodium pyruvate, 1% penicillin-streptomycin-glutamine solution, and 5 ng/ml of fibroblast growth factor-2 (FGF-2) (all from Gibco). Cells were seeded at a density of 3200 cells/cm^2^ until they reached 90% confluency. The medium was replaced twice a week.

### Chondrogenic and osteogenic differentiation

MSOD-B cells and their modified progeny were harvested from culture flasks by adding Trypsin-EDTA (0.25%) (Gibco) and subsequently seeded on cylindrical Collagen Type I scaffold (Avitene^TM^ Ultrafoam^TM^ Collagen Sponge, Davol) of 6 mm in diameter and 3 mm in thickness at a density of 2 x 10^6^ cells per scaffold in 12 well plates coated with 1 % agarose (Sigma) for chondrogenic differentiation and in a 12-well plate for osteogenic differentiation. Tissue constructs were cultured for three weeks in chondrogenic medium (DMEM supplemented with 1% penicillin-streptomycin-glutamine, 1% HEPES (1M), 1% sodium pyruvate (100 mM), 1% ITS (100x) (Insulin, Transferrin, Selenium) (all from Gibco), 0,47 mg/ml linoleic acid (Sigma), 0.12% bovine serum albumin (25 mg/mL) (BSA) (Sigma), 0.1 mM ascorbic acid (Sigma), 10^-7^ M dexamethasone (Sigma) and 10 ng/ml TGF-β3 (Novartis)). The cells in the 12 well plates were supplemented for three weeks with osteogenic (or expansion medium, control) differentiation medium (ɑ-minimum essential medium (ɑMEM) with 10% fetal bovine serum, 1% HEPES (1M), 1% sodium pyruvate (100 x 10^-3^M) and 1% penicillin-streptomycin-glutamine solution (100X), supplemented with 0.01M β-dexamethasone and 0.1M ascorbic acid). Media were replaced twice a week.

### Lyophilization

After three weeks of chondrogenic differentiation, the tissue constructs were rinsed twice with PBS 7.2 (without calcium/magnesium Gibco), snap frozen in liquid nitrogen for five minutes and then lyophilized using a freeze dryer (Labconco) (–80 °C and 0.05 mbar) overnight. Thereafter, the lyophilized tissue constructs were stored at 4 °C.

### Decellularization

After lyophilization, the tissue constructs were treated with a solution containing 1% SDS (Sigma-Aldrich) and DNase I (Sigma-Aldrich) to remove cellular material as according to Elder et al., Biomaterials 2009. The constructs were then thoroughly rinsed with PBS (7.2, without calcium/magnesium; Gibco) to eliminate residual chemicals. Following the washing step, the scaffolds were snap-frozen in liquid nitrogen for five minutes and re-lyophilized using a freeze dryer (Labconco) at –80°C and 0.05 mbar overnight. The resulting decellularized tissues were stored at 4°C until experimental use.

### Transfection of MSOD-B cells with gRNAs

MSOD-B cells were seeded at a density of 400,000 cells per well in 12 well plates to reach a minimum of 80% confluency the following day. The medium was replaced before transfection. The transfection was performed with a / ratio of 2 µL lipofectamine to 1 µg DNA. A DNA mix was prepared for each of the five gRNAs and the two plasmid controls. DNA mixtures were composed of 1 µg of plasmid, 2 µL of P3000 reagent and 50 µL of OptiMEM (Thermo-Fischer). A lipofectamine cocktail was prepared for all DNA mixtures consisting of 24 µL of lipofectamine 3000 in 400 µL of OptiMEM. Lipofectamine and the DNA mixtures were added at a 1:1 ratio. After 48h, cells were analyzed and mCherry positive clones were FACS sorted as single cells in 96-well plates using ARIAIII (BD Biosciences). Successfully expanded clones were then further characterized.

### ELISA

VEGF protein content was measured in supernatant collected from cells seeded at 570 cells/cm^2^ in T175 flask and cultured for 3 days. Content from engineered cartilage tissues was assessed following digestion in RIPA buffer. The Quantikine ELISA kit for Human VEGF-A Immunoassay from the R&D System was used according to the manufacturer instructions to determine protein concentration.

### Intracellular flow cytometry

MSOD-B and MSOD-B ΔRUNX2 cells were trypsinized, fixed and permeabilized using the Fixation/Permeabilization Kit (BD Biosciences). Following a blocking step in 10% normal donkey serum (Sigma), cells were stained with a primary antibody against RUNX2 (Rabbit anti-human, Thermo-Fischer PA5-82787) for one hour at room temperature. After primary antibody incubation, cells were washed and stained with an Allophycocyanin (APC) labeled secondary antibody (Donkey anti-rabbit IgG DyLight™ 649, Biolegend 406406) for 30-45 minutes at room temperature in 2% normal donkey serum. The Control samples consisted of unstained cells (negative control) and cells incubated only with the secondary antibody (secondary Ab control). Data were recorded on a BD LSRFortessa^TM^ Cell Analyzer (BD Biosciences). FCS files were analyzed using the FlowJo^TM^ software (FlowJo LLC, 10.5.3, BD Biosciences).

### Western blotting

MSOD-B and MSOD-B ΔRUNX2 cells were trypsinized and washed with ice cold PBS twice and lysed on ice for 10 min in RIPA buffer (#10017003, Thermo Fisher Scientific) supplemented with 1× proteinase and phosphatase inhibitor cocktail (#78440, Thermo Fisher Scientific). The lysates were centrifuged at 16, 000*g* for 15 min +4 °C, and the supernatants were collected. Sample buffer (Laemmli buffer (#161-0737, BioRad) supplemented with 5% 2-mercaptoethanol, 1× proteinase and phosphatase inhibitor cocktail (#78440, Thermo Fisher Scientific) was added to the supernatant at 1:1 ratio. Samples were boiled at 95 °C for 5 minutes and stored at −80 °C or kept on ice until gel loading. Proteins were separated using Bolt gels according to manufacturer’s protocol (#NW04122, #B0002, Thermo Fisher Scientific). iBlot2 system was used to transfer the proteins on polyvinylidene fluoride membrane (PVDF) membrane according to manufacturer’s protocol (#IB24001, Thermo Fisher Scientific). PVDF membrane was washed once in 1× PBST buffer (#28352, Thermo Fisher Scientific) and blocked in 2% blocking solution (#10156414, Thermo Fisher Scientific) for 1 h at room temperature. Membranes were incubated overnight at +4 °C with primary antibodies at recommended concentrations in 1% blocking solution. Membranes were washed three times (5 min for each wash) with 1× PBST buffer, and secondary HRP-conjugated antibodies in 1% blocking solution were added to the membranes at 1:5000 concentration for 1 h incubation at room temperature. Membranes were washed three times with 1× PBST and proteins were detected by chemiluminescence according to manufacturer’s protocol (#RPN2232, Thermo Fisher Scientific). The following antibodies were used. primary antibodies: RunX– (D1L7F – 12556) from Cell Signaling Technology and Actin (612656) is from Becton Dickinson. The secondary antibodies were anti-Mouse (GENA931) from Sigma-Aldrich, anti-Rabbit (NA9340V) from Thermo Fisher Scientific.

### Biochemical analysis

Lyophilized tissue constructs were digested by overnight incubation in 0.5 ml of Proteinase K solution (1 mg/mL Proteinase K, Sigma; 10 ug/ml pepstatin A, Sigma; 1 mM EDTA, Sigma; 100 mM Iodoacetamide; 50mM Tris) at pH 7.6 and 56 °C. The GAG content of digested samples was analyzed using Glycosaminoglycan Assay Blyscan kit (Biocolor) following the manufacturer instruction. DNA residues were quantified using the CyQuant NF Cell Proliferation Assay Kit (Thermo-Fisher, USA) following the manufacturer’s instructions, with an excitation wavelength of 485 nm and an emission wavelength of 535 nm.

### Mice

FoxN1 KO BALB/C (nude mice) 6–8-weeks old, were obtained from Charles River Laboratories. All mouse experiments and animal care were performed in accordance with the Lund University Animal Ethical Committee guidelines (ethical permit #M15485–18). Mice were housed at a 12-hour light cycle in individually ventilated cages at a positive air pressure and constant temperature. Mice were fed with autoclaved diet and water ad libitum. During the implantation procedure, anaesthesia was performed with 2-3 % Isoflurane (Attane). The mice were kept on a heating pad during the procedure to avoid the heat loss.

### Micro-CT scanning

In vivo samples were explanted and fixed overnight with 4% formaldehyde before ex vivo micro-CT analysis using a U-CT system (MILABS, Netherland) equipped with a tungsten x-ray source at 50 kV and 0.21 mA. Volumes were reconstituted at 10 µm isotropic voxel size. For total volume (TV) analysis, each sample was assessed with Blender (v2.82a, Netherland). Briefly, a mesh was created surrounding the 3D reconstruction of each sample and the volume occupied was then quantified. For bone volume (BV) analysis, the highly mineralized tissue volume was quantified using Seg3D (v2.2.1, NIH, NCRR, Science Computing and Imaging Institute (SCI))

### Sample preparation for histological analysis

In vitro samples were directly fixed prior to sectioning. In vivo samples were fixed and subsequently decalcified with a 10% EDTA solution, pH = 8, at 4 °C for two weeks prior to tissue embedding.

Paraffin embedding was performed on samples fixed in 4% formalin (Solveca AB). Tissues were dehydrated by immersion in consecutive solutions of 35%, 70%, 95%, and 99.5% graded ethanol solution (Solveco AB). Immersion in a 99.5% ethanol/xylene solution (1:1, Fisher Scientific) was then performed for ten minutes followed by two rinsing in xylene (Fisher Scientific) for 20 minutes. Subsequently, tissues were embedded in paraffin at 56 °C overnight, before sectioning with a microtome (Microm HM 355 Rotary Microto–e) in 5 – 10 μm sections. The sections were then dried overnight at 37 °C before staining. Prior to staining, sections were deparaffinized by two washes in xylene for 7 minutes and once in 99.5% ethanol/xylene solution (1:1) for 3 minutes. Afterwards, sections were hydrated twice in consecutive solutions of 99.5%, 95%, 70%, and 35% ethanol, for 7 minutes each.

Agarose embedding was performed on samples fixed overnight with 4% paraformaldehyde (Thermo Scientific), using 4% low-melting agarose (Sigma). Sections of 100 µm thickness were obtained using a 7000smz vibratome (Campden) with stainless steel or ceramic blades.

### Safranin O Staining

Paraffin-embedded sections were stained using Mayer’s hematoxylin solution (Sigma-Aldrich) for 10 minutes. Samples were then placed under running distilled water to remove superfluous staining from the sections. Subsequently, sections were stained with 0.01% fast green solution (Fisher Scientific) for five minutes and rinsed with 1% acetic acid solution (glacial, Fisher Scientific) for 15 seconds. After that, the slides were stained with 0.1% safranin O (Fisher Scientific) solution for 5 minutes. Dehydration and clearing were performed by immersion in 95%, 99.5% ethanol, 99.5% ethanol/xylene solution (1:1) and xylene twice successively for 2 minutes. Finally, the stained slides were mounted with glass slides using PERTEX mounting medium (PERTEX, HistoLab).

### Masson’s trichrome staining

Masson’s trichrome staining was performed using the trichrome staining kit (Sigma-Aldrich Sweden AB) according to the manufacturer’s guidelines. Briefly, tissue sections were deparaffinized and immersed in cold running deionized water for 3 minutes. Then the sections were kept in Bouin’s solution at room temperature overnight or at 56 °C for 15 minutes. The slides were washed by running tap water and stained using working Weigert’s iron hematoxylin solution (Sigma) for five minutes for nuclei detection (in black). After washing the slides, the cytoplasm was stained in red with Biebrich Scarlet-Acid fuchsin for five minutes followed by clearing the slides by immersion in working phosphotungstic/ phosphomolybdic acid solution for five minutes. Collagen stained blue by immersion in aniline blue solution for five minutes followed by clearing in 1% acetic acid (glacial, Fisher Scientific) solution diluted in distilled water (glacial, Fisher Scientific) for two minutes and washing with running deionized water. Finally, the sections were dehydrated in graded ethanol solutions (95% once, 100% twice) for two minutes each before washing with xylene twice for two minutes and mounted with PERTEX mounting medium.

### Hematoxylin and Eosin staining

Paraffin-embedded sections were stained using Mayer’s hematoxylin solution (Sigma-Aldrich) for 10 minutes. Samples were then placed under running distilled water to remove superfluous staining from the sections. The slides were immersed precisely in 1X phosphate-buffered saline (PBS) for 1 minute to intensify the blue nuclei staining while preserving tissue integrity. Following this, a thorough rinse with 3 changes of distilled water effectively removed any residual PBS, preparing the sections for the subsequent counterstaining step. Subsequently, Alcoholic-Eosin (Sigma-Aldrich) was applied for 1 minute as the counterstain without any rinsing post-application, ensuring optimal interaction of the staining solution with the tis. Dehydration and clearing were performed by immersion in 95%, 99.5% ethanol, 99.5% ethanol/xylene solution (1:1) and xylene twice successively for 2 minutes. Finally, the stained slides were mounted with glass coverslips using PERTEX mounting medium (PERTEX, HistoLab).

### Quantitative PCR

Total RNA isolation was conducted from in vitro engineered tissues utilizing the Quick-RNA extraction kit (Zymo Research, R1055) following the manufacturer’s protocol. Subsequently, cDNA extraction was carried out using cDNA synthesis kit (Invitrogen 11917020). The quantitative real-time polymerase chain reaction (qRT-PCR) was performed, utilizing the assay on demand from Applied Biosystems to assess the expression of specific genes. These genes include glyceraldehyde 3-phosphate dehydrogenase (GAPDH, HS02786624_G1), runt-related transcription factor 2 (Runx2, Hs00231692_m1), alkaline phosphatase (ALP, Hs01029144_m1), and collagen type X (ColX, Hs00166657_m1).

### Rat osteochondral defect model

Ten-to 12-week-old male Sprague Dawley rats (n=11) were purchased from JANVIER LABS (France). After 7 days of acclimation the rats were anesthetized using 3% isoflurane. Then, animals were placed on a 37°C warm heating pad in prone position. Once anesthetized, isoflurane was lowered to 2 to 2.5%, and buprenorphine (Temgesic, 30 μg/kg; Indivior Europe Ltd., Dublin, Ireland) was injected subcutaneously for analgesia. The right hind limb of the animal was shaved and carefully disinfected, and an incision was made along the skin and soft tissue to expose the right knee. After cleaning the knee laterally from soft tissue, patella was displaced laterally to expose the distal femur. An articular cartilage defect with 2mm depth and 1mm diameter was created at the trochlear groove of femur. The defect was then push-fitted with the grafts, and not secured with any sutures. The wound was closed in a layered fashion using resorbable sutures (Vicryl 45-0, Ethicon, Somerville, USA) by closing the joint capsule, followed by the muscle tissue (continuous interlaced suture), followed by closing the skin (Donati suture). Animals started load bearing immediately after surgery. After 3 weeks (n=11) animals were anesthetized by isoflurane inhalation (3%), followed by CO2 asphyxiation, and right-femur were harvested prior to subsequent micro-CT and histological characterization.

### Histology grading method

Histological sections from rat femur condyles were stained with Safranin-O, Masson’s trichrome, and H&E staining. For each sample, sections were compiled at different depth (top, middle and bottom) defined to encompass the full tissue characteristics. A semi-quantitative analysis of the repaired tissue was performed by utilizing a customized adaptation of the histological grading system initially outlined by Wakitani et al^66^. This modified scale encompasses six distinct parameters: cellular morphology, matrix staining, surface regularity, cartilage thickness, regenerated subchondral bone formation, and integration with neighbouring cartilage. Each parameter was graded on a numerical scale ranging from 0 to 4 points by independent expert observers, where a maximum total score of 16 indicates the presence of tissue displaying characteristics akin to entirely healthy and normal tissue (Table 1).

### Cartilage quantification method

An evaluation of three distinct sites within the condyle defect was conducted to ascertain the proportion of cartilage in each knee. Utilizing the measured femoral condyle thickness (200μm) and defect length (1mm), a standardized Region of Interest (ROI) was defined using ImageJ. The percentage of cartilage-positive regions within each defect was calculated in relation to the total ROI size, offering a quantitative assessment of cartilage regeneration. This identical approach was applied to evaluate the percentage of cartilage in the healthy control samples.

### Trabecular thickness (Tb.Th) and trabecular separation (Tb.Sp)

Image analysis was performed by ImageJ. To evaluate the defect space, a rectangular interface region (2 × 1 mm) was defined as the Region of Interest. Trabecular thickness (Tb.Th) and trabecular separation (Tb.Sp),was calculated inside the ROI using BoneJ plugin.

### Chick chorioallantoic membrane (CAM) assay

Fertilized eggs from Lohmann Brown chicken were commercially purchased and incubated 6 days prior (Day –6) in a BINDER incubator at 37.5 °C with constant humidity. A small window in the shell was opened 3 days prior (Day –3) to the start of the experiment under aseptic conditions. The window was resealed with adhesive tape and eggs were returned to the incubator. On day 0, MSOD-B and MSOD-BΔV1 in vitro differentiated and lyophilized samples were placed on top of the CAM. Eggs were resealed and returned to the incubator. Pictures were taken with a brightfield microscope (LEICA S9i) on Day 4 and analysed for vascular density.

### Vascular density quantification

To assess vascular density, the Quantitative Vascular Analysis Tool (Q-VAT)^30^ was utilized. The images were acquired using a digital microscope. These images were then segmented into tiles for detailed analysis. The binary vascular masks were generated using the Q-VAT’s automated capabilities, to distinguish vascular structures within the tissue. The tool then facilitated the separation of vessel measurements based on their diameters, allowing for a differentiated quantification of macro– and microvasculature. These binary masks were then processed to calculate the vascular density, which refers to the proportion of the tissue area occupied by vessels. This quantitative metric was evaluated for each tile, and the mean vascular density was computed across all tiles for each tissue sample. As part of the analysis, we applied Q-VAT to double or triple-stained slides to quantify the overlapping percentage of vessels, comparing various time points to observe changes over the course of the experiment.

### Immunofluorescence

Agarose-embedded sections were treated with 0.5% Triton X-100 (Sigma) in PBS supplemented with 20% donkey serum (Jackson ImmunoResearch) for one hour at room temperature (RT) for blocking and permeabilization. After blocking/permeabilization, sections were stained with primary antibodies (mouse anti-Collagen II (Invitrogen, MA137493), Anti-Collagen 1 (Abcam, ab138492), VEGF polyclonal antibody (Bioss Antibodies, bs-0279R), rabbit anti-Collagen Type X (ColX) (abbexa, abx101469), Goat anti mouse/rat CD31 (R&D system, YZU0119121), VEGF at 4 °C overnight. Sections were washed three times using cold PBS with 0.1% Triton X-100, 20 minutes each, followed by staining with secondary antibodies (CF568 donkey anti-rabbit (Biotium, 20098), CF567 anti-goat (Invitrogen, A11057), CF633 donkey anti-mouse (Sigma-Aldrich, SAB4600131 in 2% donkey serum for three hours at room temperature. Sections were washed three times using cold PBS with 0.1% Triton X-100, 20 minutes each once more.

All sections were mounted using the Vectashield antifade mounting medium containing DAPI (Vector Laboratories, H1200). A LSM780 confocal microscope (Zeiss) with a 10x/20x objective was used to capture images and image series of whole tissue sections (z-step = 2 µm). The IMARIS 9.5 software package (Oxford Instruments) was used to analyze the data.

## Acknowledgements

We express our gratitude to the following facilities and individuals for their invaluable support and contributions to this study: FACS Facility (Anna Hammarberg), Cell and Gene Therapy core facility (Pia Johansson) and Imaging Facility at Lund Stem Cell Centre, as well as the Lund University Bioimaging Centre, for providing access to the X-ray CT system. This research study was made possible by financial support through the Knut and Alice Wallenberg grant, the Medical Faculty at Lund University, the Swedish Research Council (Starting grant to P.E.B), the European Research Council (Starting grant hOssicle #948588 to P.E.B), and Region Skåne. Figures were generated using BioRender. Special thanks to Ani Grigoryan for her expertise and valuable insights regarding FACS experiments. Christian Hansen deserves our appreciation for providing the mcherry plasmid and offering valuable guidance and knowledge related to CRISPR-Cas9.

---

**Supplementary figure 1.**
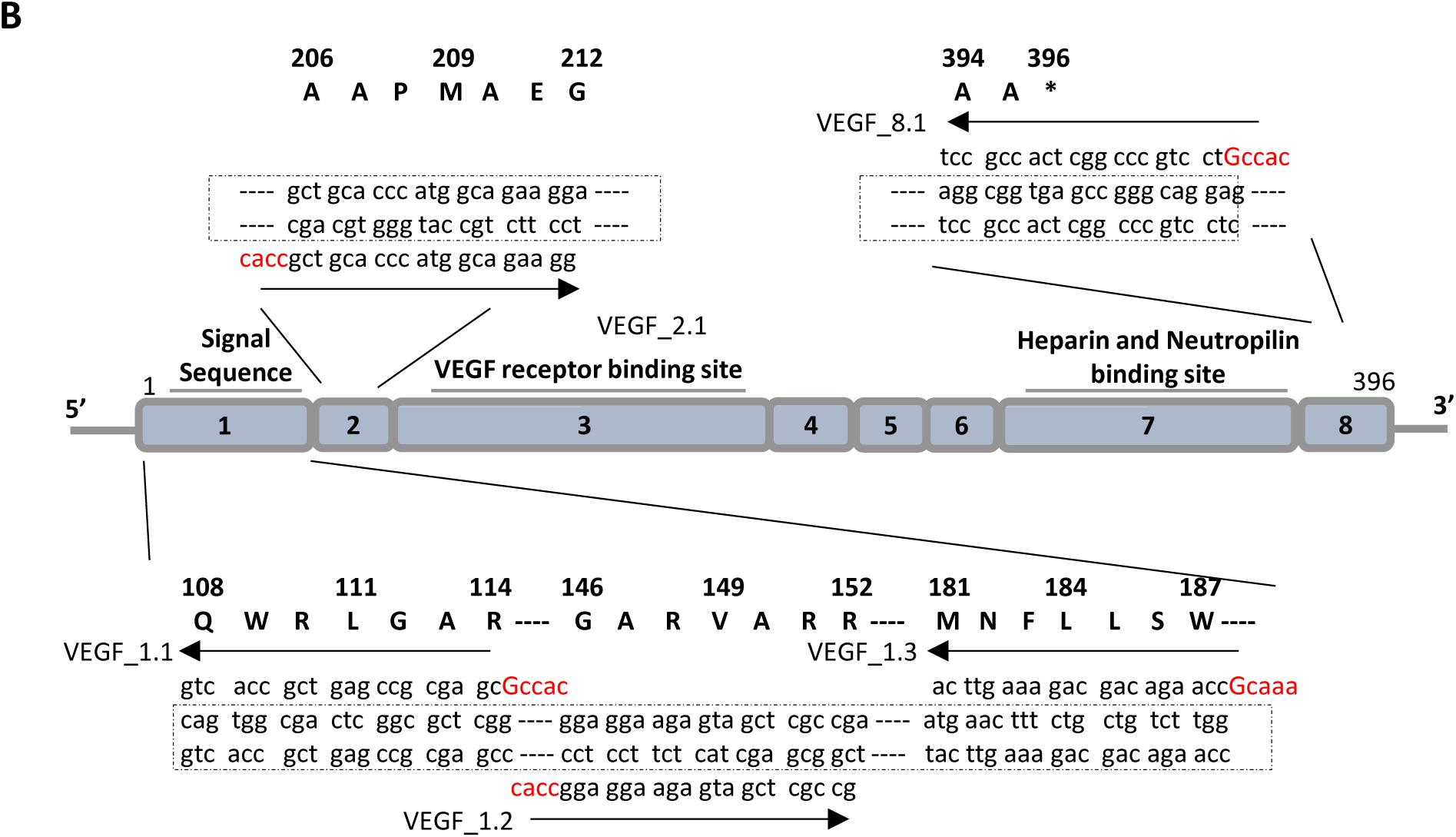
CRISPR/Cas9 editing of mesenchymal cell lines lead to efficient VEGF knockout in cartilage tissues. A) Implemented genetic elements of the Mesenchymal Sword Of Damocles BMP-2 (MSOD-B) cells consisting of the human telomerase reverse transcriptase (hTERT), an inducible caspase 9 (iCaspase) death system and the bone morphogenetic protein type-2. All systems are constitutively expressed and further described in Pigeot et al. Advanced Materials 2021. B) Overview of the human VEGF exon structure, nucleotide sequences of targeting gRNAs and their corresponding amino acid sequences.

**Supplementary figure 2.**
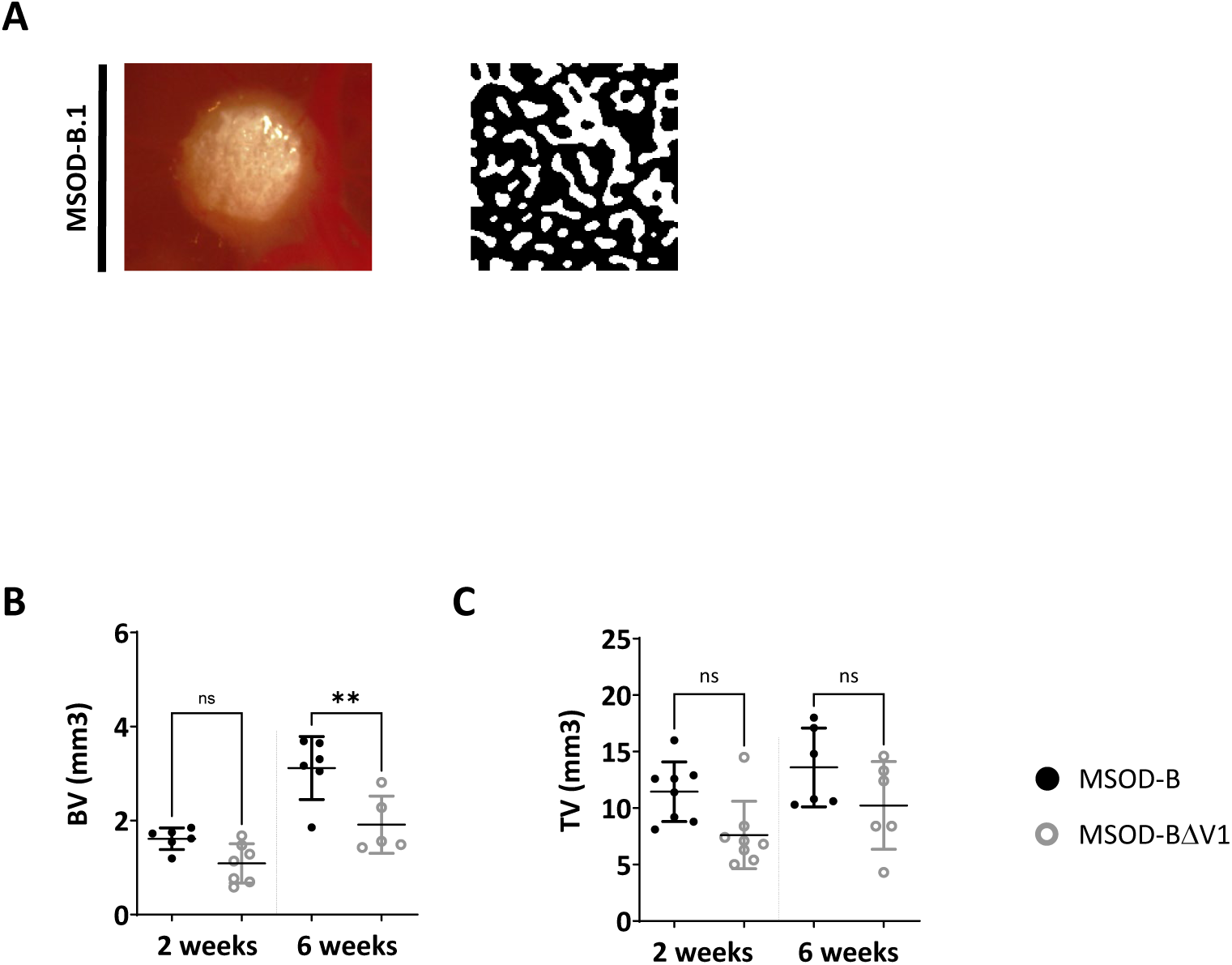
VEGF knockout cartilage tissues retain bone remodeling capacity despite reduced early-stage vascularization. A) Sequential processing of a vascular tissue section using Q-VAT, depicting the original image, binary mask creation, and vessel segmentation for quantitative vascular density analysis. B) Microtomography analysis (Bone/mineralized volume) showed significant differences between in vivo constructs at both two weeks and six weeks time points (One way ANOVA, n=3, **p < 0.01, n.s.= not significant). C) Microtomography analysis (total volume) did not show significant difference between in vivo constructs at either two weeks or six weeks time points (One way ANOVA, n=3, n.s.= not significant) (BV: Bone Volume, TV: Total Volume).

**Supplementary figure 3.**
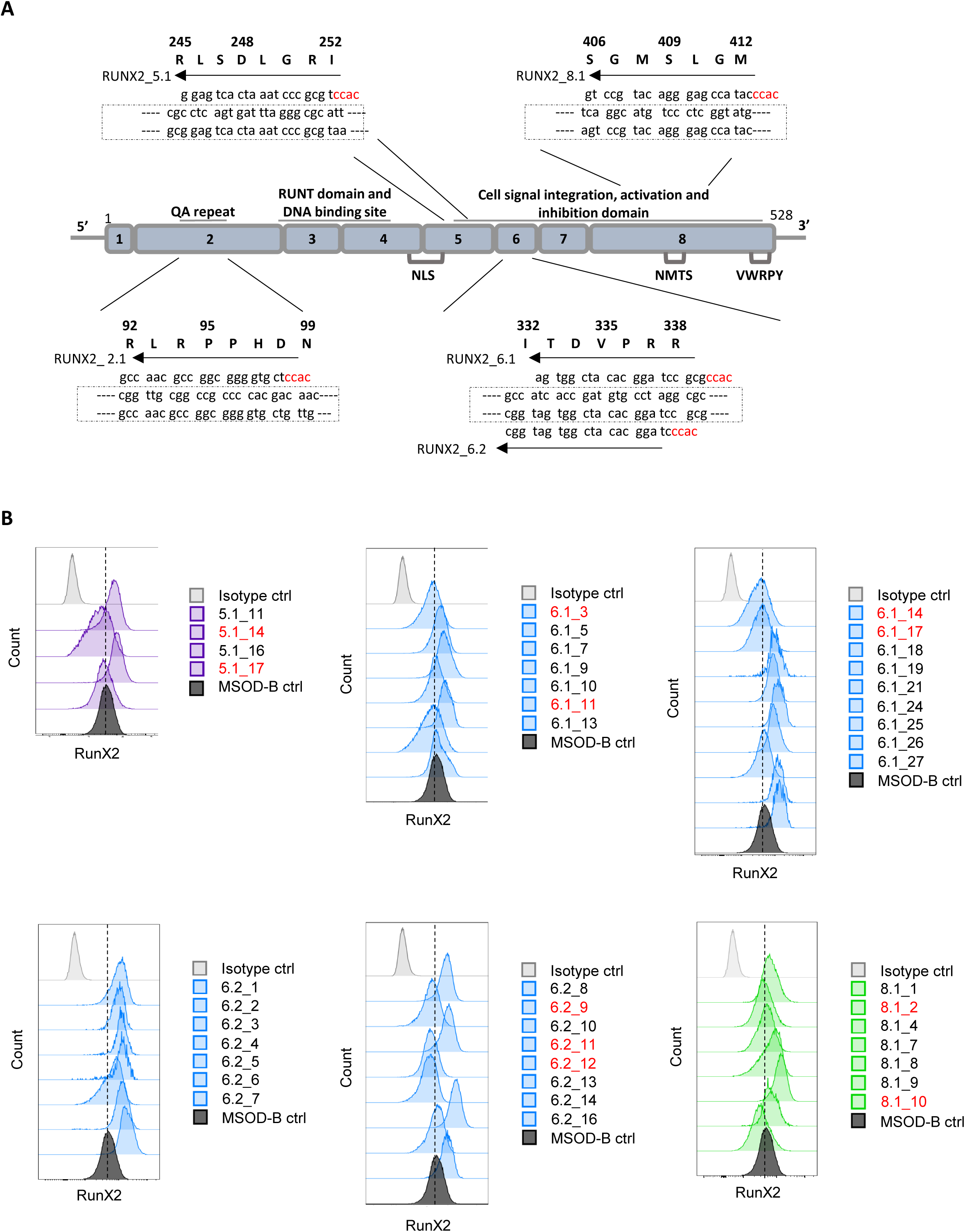
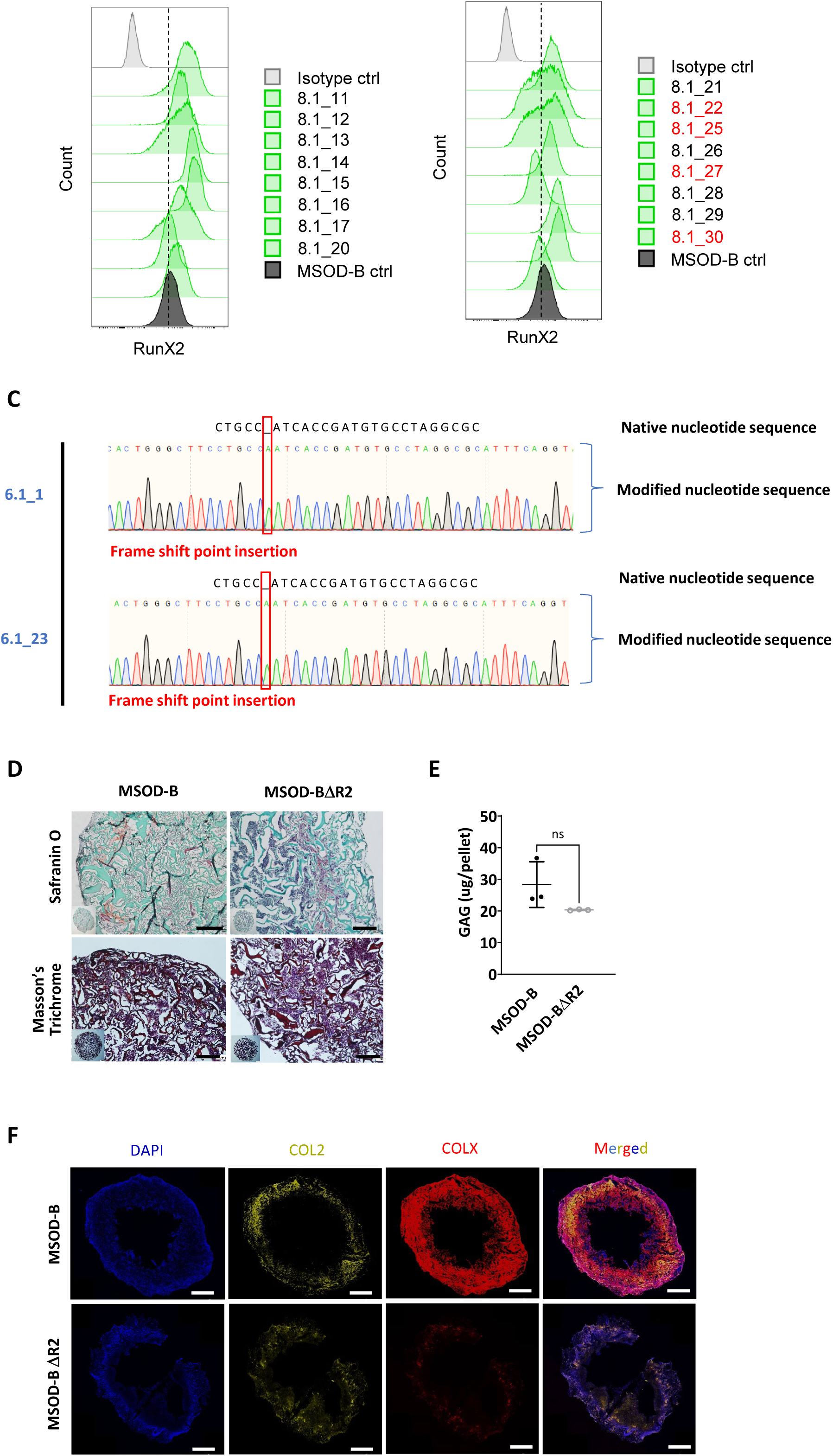
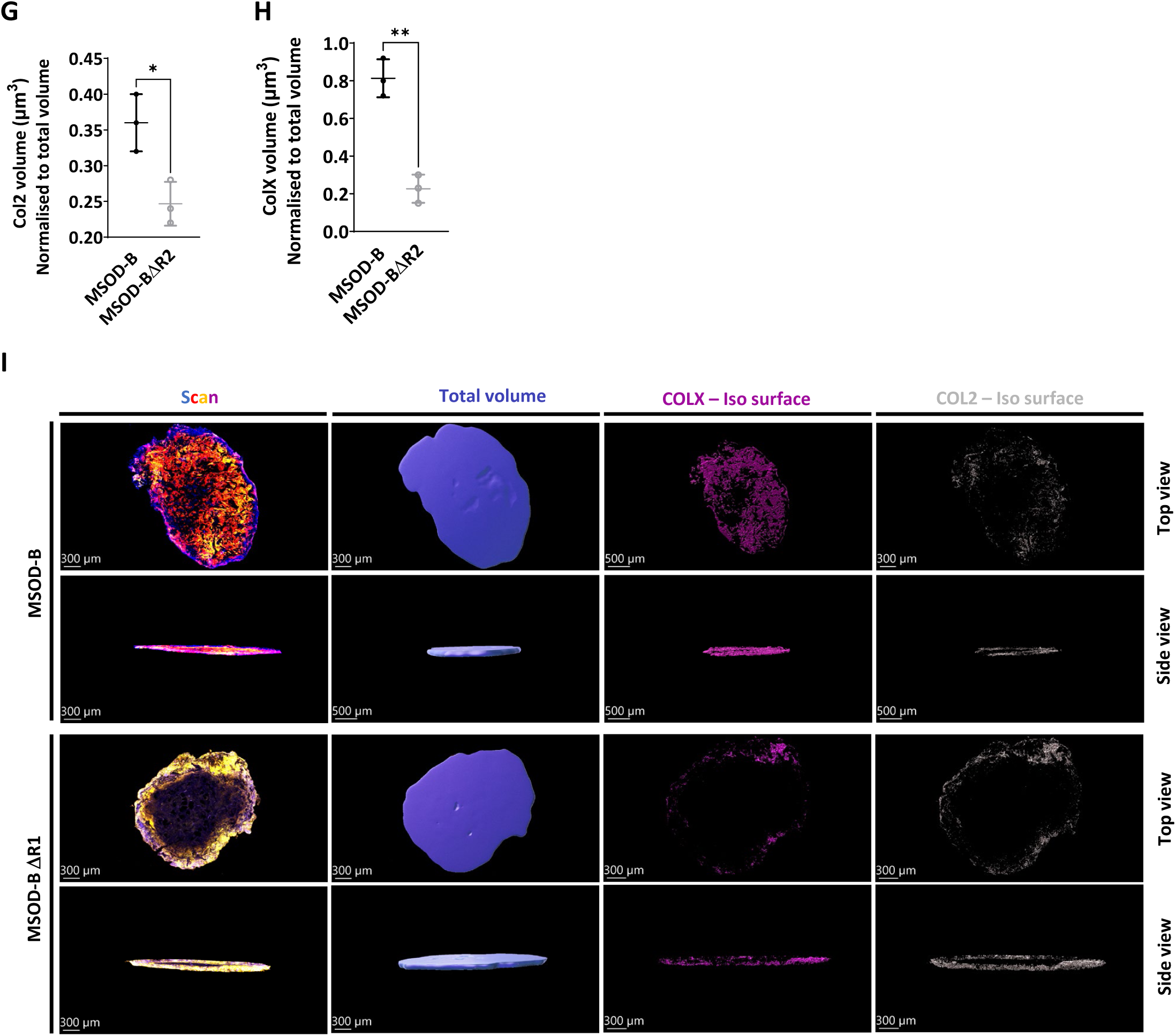
RUNX2 knockout does not prevent chondrogenic differentiation but impairs hypertrophy. A) Overview of human RUNX2 exon structure, nucleotide sequences of targeting gRNAs and its corresponding amino acid sequence. B) Intracellular flow cytometry for RUNX2 detection in MSOD-B and RUNX2-edited clones. Clones whose peaks fell left of the dotted line were selected for sequencing. C) Sanger sequencing results of DNA extracted from MSOD-BΔR samples and the modifications compared to MSOD-B DNA sequence. D) Histological analysis of in vitro constructs from using Safranin O and Masson trichrome staining displayed the presence of glycosaminoglycans (GAG) and collagen content respectively, indicating the presence of cartilage formation in MSOD-BΔR2 (Scale bars = 200 µm). E) Quantitative assessment of the total GAG content confirms the presence of cartilage formation in both constructs thus confirming RUNX2 knockout in hMSCs (MSOD-BΔR2) retain cartilage formation. (unpaired t-test, n=3, n.s. = not significant). F) Reduction of RUNX2 expression in cartilage tissues lead to reduced hypertrophy. Representative immunofluorescence staining images of COL2, COLX and DAPI tissue section of in vitro cartilage constructs shows reduction of COLX expression in knockout construct (MSOD-BΔR2) indicating reduction of hypertrophy. (Scale bars = 500 µm). G) Quantitative analysis of the immunofluorescent staining sections using IMARIS software displayed significant difference in the expression of COL2, demonstrating reduction of cartilage formation in knockout constructs (MSOD-BΔR2) (unpaired t-test, n=3, *p < 0.05). H) Quantitative analysis of the immunofluorescent staining sections using IMARIS software displayed a significant reduction in expression of COLX, confirming the disruption of hypertrophy in knockout constructs (unpaired t-test, n=3, **p < 0.01). I) Representative isosurface images of immunofluorescent staining sections constructed using IMARIS software.

**Supplementary figure 4.**
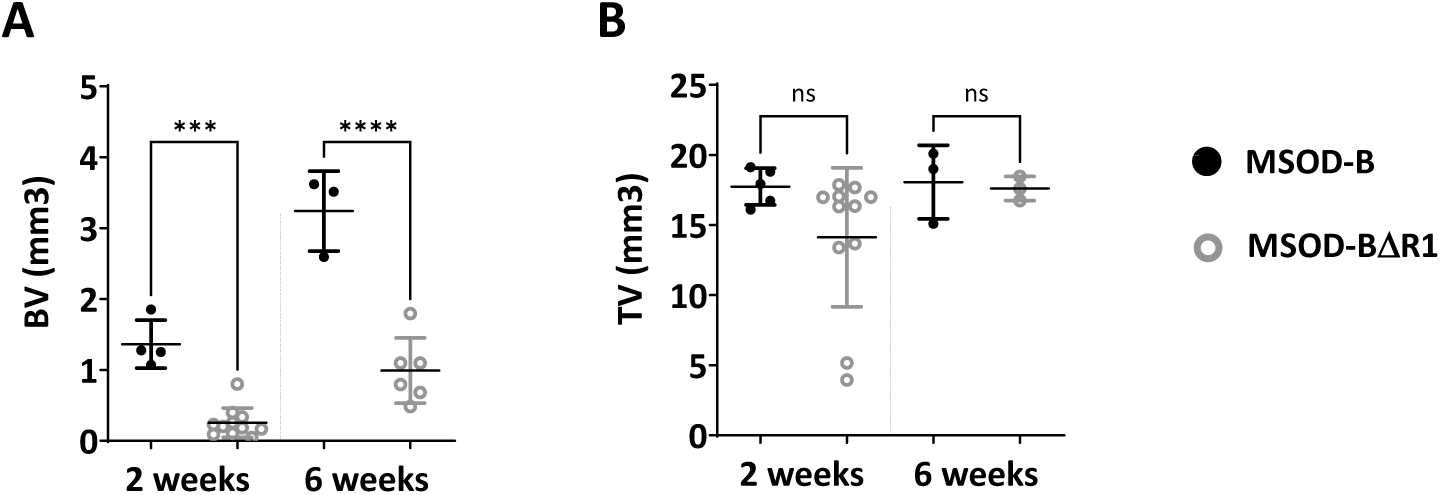
RUNX2 knockout in cartilage tissues disrupts effective bone formation. A) Microtomography analysis (Bone/mineralized volume) showed very high significant difference between in vivo constructs at both two weeks and six weeks time points (One was ANOVA, n=3, ***p < 0.001, ****p < 0.0001). B) Microtomography analysis (total volume) showed significant difference between in vivo constructs at two weeks time point but no significant difference at six weeks time point (One was ANOVA, n=3, n.s. = not significant) (BV: Bone Volume, TV: Total Volume).

**Supplementary figure 5.**
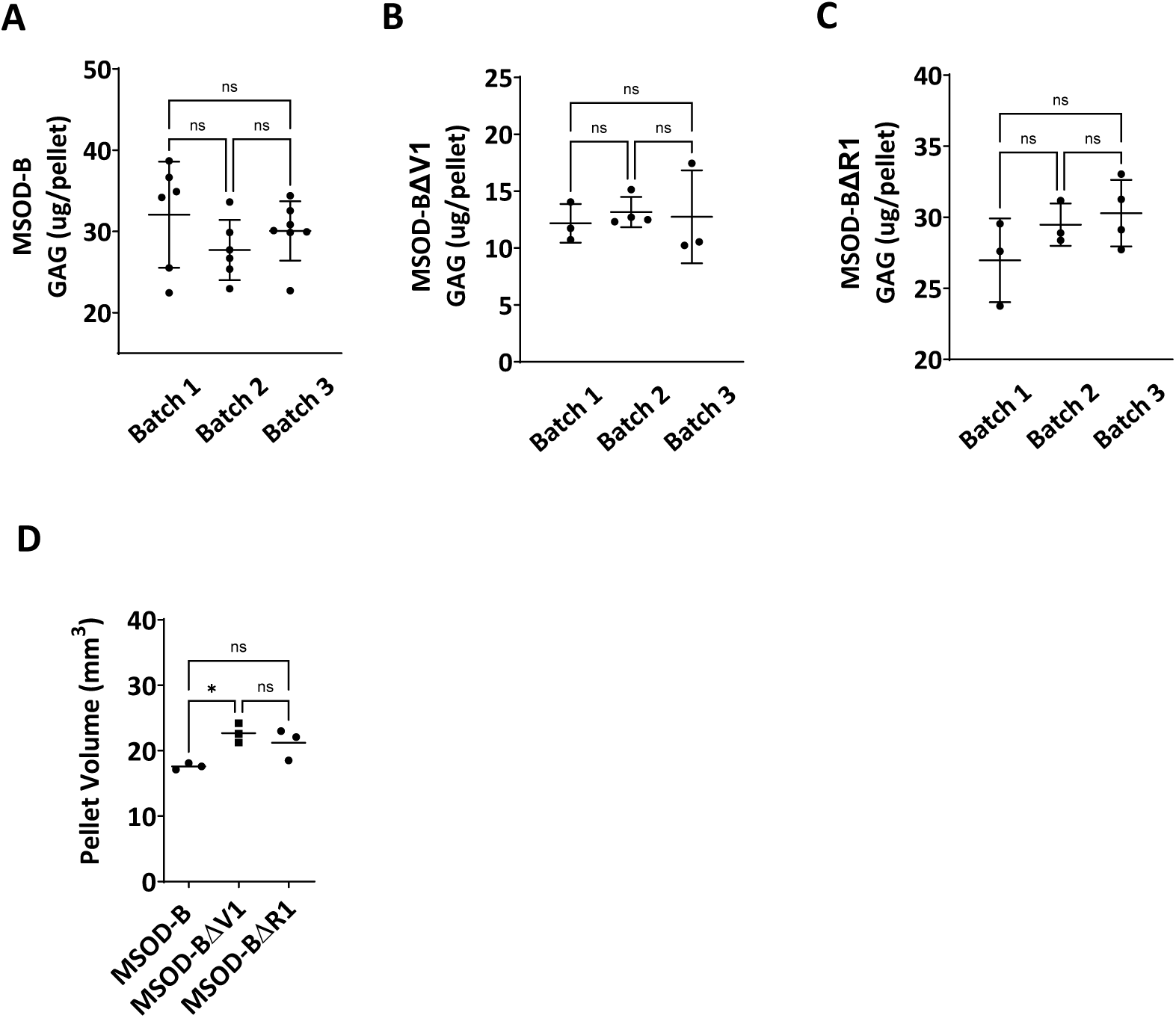
Characterization of engineered cartilage tissues post-lyophilization. A) Quantitative assessment of the total GAG content in MSOD-B in vitro differentiated constructs across three different batch productions. One way ANOVA test, n=6-7 biological replicates, n.s. = not significant. B) Quantitative assessment of the total GAG content in MSOD-BΔV1 in vitro differentiated constructs across three different batch productions. One way ANOVA test, n=3-4 biological replicates, n.s.=not significant. C) Quantitative assessment of the total GAG content in MSOD-BΔR1 in vitro differentiated constructs across three different batch productions. One way ANOVA test, n=3 biological replicates, ***p < 0.001, ****p < 0.0001, n.s =not significant. D) Quantitative assessment of cartilage tissue volume in MSOD-B, MSOD-BΔV1 and MSOD-BΔR1 tissues. One way ANOVA test, n=3 biological replicates, *p < 0.05, n.s.=not significant.

**Supplementary figure 6.**
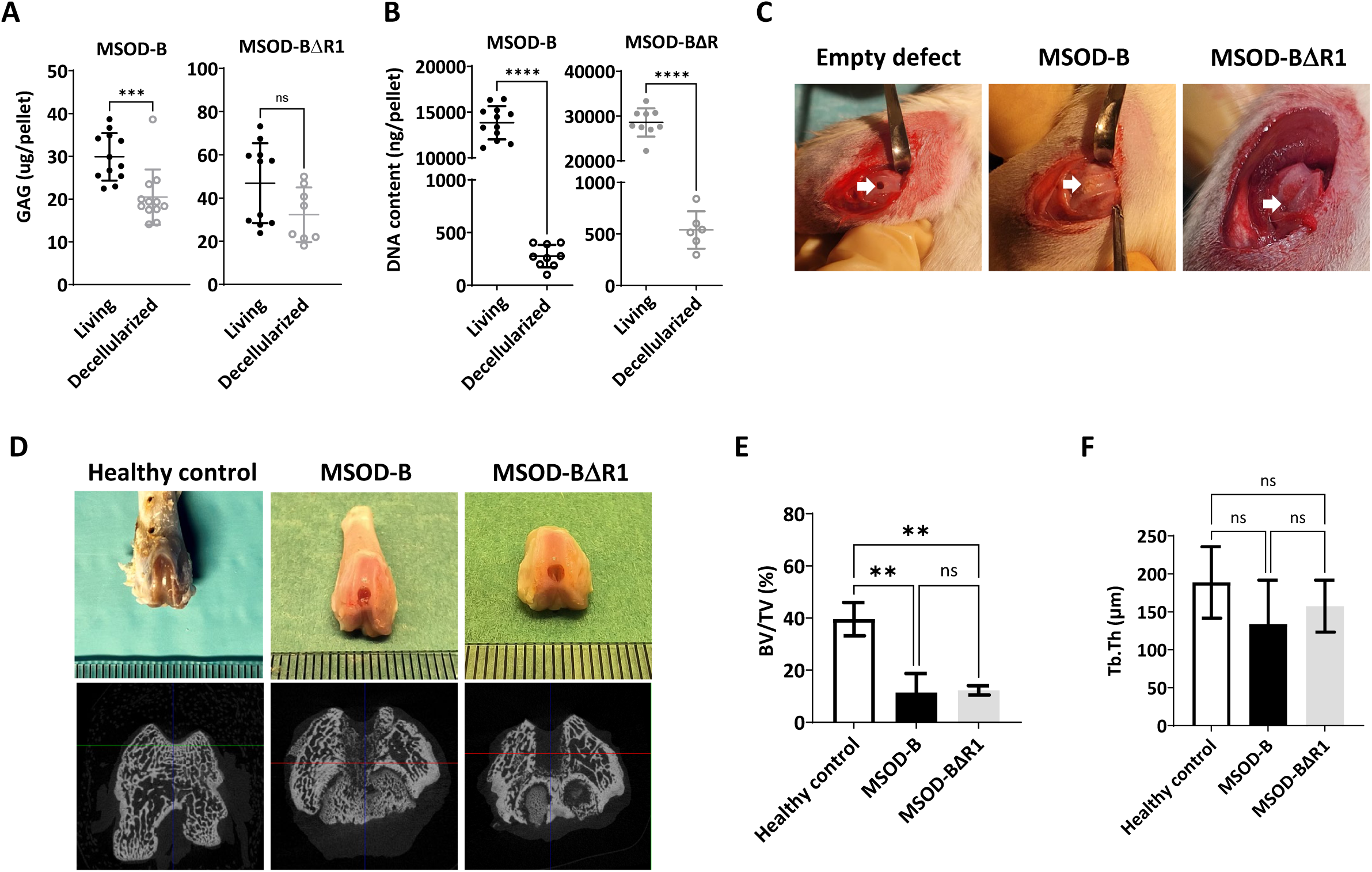
RUNX2 knockout in cartilage tissues exhibit better cartilage regeneration and maintenance in osteochondral defect in rats. A) Quantitative assessment of the total GAG content in MSOD-B and MSOD-BΔR1 in vitro differentiated constructs before (living) and after (Decellularized) the decellularization process. unpaired t-test, n=12 biological replicates, ***p < 0.001, n.s. = not significant B) DNA quantification of in vitro engineered cartilage grafts before (living) and after (Decellularized) the decellularization process. unpaired t-test, n=12 biological replicates, ****p < 0.0001. C) Macroscopic images osteochondral defect model in rats depicting in of empty defect, defect filled with MSOD-B and defect filled with MSOD-BΔR1 respectively. D) Macroscopic images of extracted rat knees depicting healthy control without any defect, defect filled with MSOD-B and defect filled with MSOD-BΔR1 respectively and microtomography images of same respective constructs. E). No significant difference in BV/TV is observed between MSOD-B & MSOD-BΔR1 samples despite significantly reduced BV/TV ratio in both samples compared to healthy control. One way ANOVA test, n=3 biological replicates, **p < 0.01. F) Image J based quantification of Trabecular thickness (tb.Th). One way ANOVA test, n=3 biological replicates, n.s = not significant.

**Supplementary table 1:** Histological grading system. The histological graded system is modified and adapted from previously established model as mentioned in materials and methods section. This modified scale incorporates six distinct parameters to comprehensively evaluate the quality of tissue repair

| Category | Point |
| --- | --- |
| **Cell morphology** |  |
| Hyaline cartilage | 4 |
| Mostly hyaline cartilage or fibrocartilage | 3 |
| Mostly fibrocartilage | 2 |
| Mostly non-cartilage | 1 |
| Non-cartilage | 0 |
| **Matrix-staining (metachromasia)** |  |
| Normal | 3 |
| Slightly reduced | 2 |
| Markedly reduced | 1 |
| No metachromatic staining | 0 |
| **Surface regularity<sup>a</sup>** |  |
| Smooth | 3 |
| Moderate | 2 |
| Irregular | 1 |
| Severely irregular | 0 |
| **Thickness of cartilage (%)<sup>b</sup>** |  |
| 121–150 | 1 |
| 81–120 | 2 |
| 51–80 | 1 |
| 0–50 | 0 |
| **Regenerated subchondral bone** |  |
| Good | 2 |
| Moderate | 1 |
| Poor | 0 |
| **Integration with adjacent cartilage** |  |
| Both edges integrated | 2 |
| One edge integrated | 1 |
| Neither edge integrated | 0 |
| **Total maximum** | **16** |

•<sup>a</sup> Total smooth area of the reparative cartilage compared with the entire area of the cartilage defect.
•<sup>b</sup> Average thickness of the reparative cartilage compared with that of the surrounding cartilage.

## Notes

### Competing Interest Statement

The authors have declared no competing interest.

### Summary of Updates

Text modifications and addition of new results.

## References

1. Bonnans C, Chou J, Werb Z. Remodelling the extracellular matrix in development and disease. Nat Rev Mol Cell Biol. 2014;15(12):786–801. doi:10.1038/nrm3904

2. Hussey GS, Dziki JL, Badylak SF. Extracellular matrix-based materials for regenerative medicine. Nat Rev Mater. 2018;3(7):159–173. doi:10.1038/s41578-018-0023-x

3. Fattahi R, Chamkhorami FM, Taghipour N, Keshel SH. The effect of extracellular matrix remodeling on material-based strategies for bone regeneration: Review article. Tissue Cell. 2022;76(January):101748. doi:10.1016/j.tice.2022.101748

4. Mouw JK, Ou G, Weaver VM. Extracellular matrix assembly: A multiscale deconstruction. Nat Rev Mol Cell Biol. 2014;15(12):771–785. doi:10.1038/nrm3902

5. Assunção M, Dehghan-Baniani D, Yiu CHK, Später T, Beyer S, Blocki A. Cell-Derived Extracellular Matrix for Tissue Engineering and Regenerative Medicine. Front Bioeng Biotechnol. 2020;8(December):1–10. doi:10.3389/fbioe.2020.602009

6. Hinderer S, Layland SL, Schenke-Layland K. ECM and ECM-like materials – Biomaterials for applications in regenerative medicine and cancer therapy. Adv Drug Deliv Rev. 2016;97:260–269. doi:10.1016/j.addr.2015.11.019

7. Hinderer S, Layland SL, Schenke-Layland K. ECM and ECM-like materials – Biomaterials for applications in regenerative medicine and cancer therapy. Adv Drug Deliv Rev. 2016;97:260–269. doi:10.1016/j.addr.2015.11.019

8. Fattahi R, Chamkhorami FM, Taghipour N, Keshel SH. The effect of extracellular matrix remodeling on material-based strategies for bone regeneration: Review article. Tissue Cell. 2022;76(January):101748. doi:10.1016/j.tice.2022.101748

9. Tang G, Liu Z, Liu Y, et al. Recent Trends in the Development of Bone Regenerative Biomaterials. Front Cell Dev Biol. 2021;9(May):1–18. doi:10.3389/fcell.2021.665813

10. Gao C, Peng S, Feng P, Shuai C. Bone biomaterials and interactions with stem cells. Bone Res. 2017;5(May):1–33. doi:10.1038/boneres.2017.59

11. Filippi M, Born G, Chaaban M, Scherberich A. Natural Polymeric Scaffolds in Bone Regeneration. Front Bioeng Biotechnol. 2020;8(May). doi:10.3389/fbioe.2020.00474

12. Reddy MSB, Ponnamma D, Choudhary R, Sadasivuni KK. A comparative review of natural and synthetic biopolymer composite scaffolds. Polymers (Basel). 2021;13(7). doi:10.3390/polym13071105

13. Decaris ML, Leach JK. Design of experiments approach to engineer cell-secreted matrices for directing osteogenic differentiation. Ann Biomed Eng. 2011;39(4):1174–1185. doi:10.1007/s10439-010-0217-x

14. Harvestine JN, Orbay H, Chen JY, Sahar DE, Leach JK. Cell-secreted extracellular matrix, independent of cell source, promotes the osteogenic differentiation of human stromal vascular fraction. J Mater Chem B. 2018;6(24):4104–4115. doi:10.1039/c7tb02787g

15. Hoch AI, Mittal V, Mitra D, Vollmer N, Zikry CA, Leach JK. Cell-secreted matrices perpetuate the bone-forming phenotype of differentiated mesenchymal stem cells. Biomaterials. 2016;74:178–187. doi:10.1016/j.biomaterials.2015.10.003

16. Haumer A, Bourgine PE, Occhetta P, Born G, Tasso R, Martin I. Delivery of cellular factors to regulate bone healing. Adv Drug Deliv Rev. 2018;129:285–294. doi:10.1016/j.addr.2018.01.010

17. Sadr N, Pippenger BE, Scherberich A, et al. Enhancing the biological performance of synthetic polymeric materials by decoration with engineered, decellularized extracellular matrix. Biomaterials. 2012;33(20):5085–5093. doi:10.1016/j.biomaterials.2012.03.082

18. Prewitz MC, Seib FP, von Bonin M, et al. Tightly anchored tissue-mimetic matrices as instructive stem cell microenvironments. Nat Methods. 2013;10(8):788–794. doi:10.1038/nmeth.2523

19. Bourgine PE, Gaudiello E, Pippenger B, et al. Engineered Extracellular Matrices as Biomaterials of Tunable Composition and Function. Adv Funct Mater. Published online 2017. doi:10.1002/adfm.201605486

20. Pigeot S, Klein T, Gullotta F, et al. Manufacturing of Human Tissues as off-the-Shelf Grafts Programmed to Induce Regeneration. Advanced Materials. 2021;33(43). doi:10.1002/adma.202103737

21. Grigoryan A, Zacharaki D, Balhuizen A, et al. Engineering of human mini-bones for the standardized modeling of healthy hematopoiesis, leukemia and solid tumor metastasis. bioRxiv. Published online 2021:2021.09.11.459806. https://www.biorxiv.org/content/10.1101/2021.09.11.459806v1%0Ahttps://www.biorxiv.org/content/10.1101/2021.09.11.459806v1.abstract

22. Golchin A, Shams F, Karami F. Advancing Mesenchymal Stem Cell Therapy with CRISPR/Cas9 for Clinical Trial Studies. In: Cell Biology and Translational Medicine,. Vol 8.; 2019:89–100. doi:10.1007/5584_2019_459

23. Hazrati A, Malekpour K, Soudi S, Hashemi SM. CRISPR/Cas9-engineered mesenchymal stromal/stem cells and their extracellular vesicles: A new approach to overcoming cell therapy limitations. Biomedicine & Pharmacotherapy. 2022;156:113943. doi:10.1016/j.biopha.2022.113943

24. Geurts MH, Clevers H. CRISPR engineering in adult stem cell-derived organoids for gene repair and disease modelling. 2023;1(January):32–45. doi:10.1038/s44222-022-00013-5

25. Hu K, Olsen BR. The roles of vascular endothelial growth factor in bone repair and regeneration. Bone. 2016;91:30–38. doi:10.1016/j.bone.2016.06.013

26. Duda GN, Geissler S, Checa S, Tsitsilonis S, Petersen A, Schmidt-Bleek K. The decisive early phase of bone regeneration. Nat Rev Rheumatol. 2023;19(February):78–95. doi:10.1038/s41584-022-00887-0

27. Ran FA, Hsu PD, Wright J, Agarwala V, Scott DA, Zhang F. Genome engineering using the CRISPR-Cas9 system. Nat Protoc. 2013;8(11):2281–2308. doi:10.1038/nprot.2013.143

28. joces_113_1_59.

29. Wang L, Wan L, Zhang T, et al. A Combined Treatment of BMP2 and Soluble VEGFR1 for the Enhancement of Tendon-Bone Healing by Regulating Injury-Activated Skeletal Stem Cell Lineage. American Journal of Sports Medicine. 2024;52(3):779–790. doi:10.1177/03635465231225244

30. Callewaert B, Gsell W, Himmelreich U, Jones EAV. Q-VAT: Quantitative Vascular Analysis Tool. Front Cardiovasc Med. 2023;10. doi:10.3389/fcvm.2023.1147462

31. Pigeot S, Klein T, Gullotta F, et al. Manufacturing of Human Tissues as off-the-Shelf Grafts Programmed to Induce Regeneration. Advanced Materials. 2021;33(43). doi:10.1002/adma.202103737

32. Otto F, Thornell AP, Crompton T, et al. Cbfa1, a candidate gene for cleidocranial dysplasia syndrome, is essential for osteoblast differentiation and bone development. Cell. 1997;89(5):765–771. doi:10.1016/S0092-8674(00)80259-7

33. Komori T, Yagi * H., Nomura S, Yamaguchi A. Targeted disruption of. Proc Natl Acad Sci U S A. 1998;95(15):8692–8697.

34. Kim HJ, Kim WJ, Ryoo HM. Post-Translational Regulations of Transcriptional Activity of RUNX2. Mol Cells. 2020;43(2):160–167. doi:10.14348/molcells.2019.0247

35. Hattori EY, Masuda T, Mineharu Y, et al. A RUNX-targeted gene switch-off approach modulates the BIRC5/PIF1-p21 pathway and reduces glioblastoma growth in mice. Commun Biol. 2022;5(1):1–11. doi:10.1038/s42003-022-03917-5

36. Chen H, Ghori-Javed FY, Rashid H, et al. Runx2 regulates endochondral ossification through control of chondrocyte proliferation and differentiation. Journal of Bone and Mineral Research. 2014;29(12):2653–2665. doi:10.1002/jbmr.2287

37. Elder BD, Eleswarapu S V., Athanasiou KA. Extraction techniques for the decellularization of tissue engineered articular cartilage constructs. Biomaterials. 2009;30(22):3749–3756. doi:10.1016/j.biomaterials.2009.03.050

38. Komori T. Runx2, an inducer of osteoblast and chondrocyte differentiation. Histochem Cell Biol. 2018;149(4):313–323. doi:10.1007/s00418-018-1640-6

39. Maehara H, Sotome S, Yoshii T, et al. Repair of large osteochondral defects in rabbits using porous hydroxyapatite/collagen (HAp/Col) and fibroblast growth factor-2 (FGF-2). Journal of Orthopaedic Research. 2010;28(5):677–686. doi:10.1002/jor.21032

40. Kusumbe AP, Ramasamy SK, Adams RH. Coupling of angiogenesis and osteogenesis by a specific vessel subtype in bone. Nature. 2014;507(7492):323–328. doi:10.1038/nature13145

41. Tuckermann J, Adams RH. The endothelium–bone axis in development, homeostasis and bone and joint disease. Nat Rev Rheumatol. 2021;17(10):608–620. doi:10.1038/s41584-021-00682-3

42. Largo RD, Burger MG, Harschnitz O, et al. VEGF Over-Expression by Engineered BMSC Accelerates Functional Perfusion, Improving Tissue Density and In-Growth in Clinical-Size Osteogenic Grafts. Front Bioeng Biotechnol. 2020;8(July):1–13. doi:10.3389/fbioe.2020.00755

43. Yang YQ, Tan YY, Wong R, Wenden A, Zhang LK, Rabie ABM. The role of vascular endothelial growth factor in ossification. Int J Oral Sci. 2012;4(2):64–68. doi:10.1038/ijos.2012.33

44. Burger MG, Grosso A, Briquez PS, et al. Robust coupling of angiogenesis and osteogenesis by VEGF-decorated matrices for bone regeneration. Acta Biomater. 2022;149:111–125. doi:10.1016/j.actbio.2022.07.014

45. Behr B, Tang C, Germann G, Longaker MT, Quarto N. Locally applied vascular endothelial growth factor A increases the osteogenic healing capacity of human adipose-derived stem cells by promoting osteogenic and endothelial differentiation. Stem Cells. 2011;29(2):286–296. doi:10.1002/stem.581

46. Zuo WH, Zeng P, Chen X, Lu YJ, Li A, Wu J Bin. Promotive effects of bone morphogenetic protein 2 on angiogenesis in hepatocarcinoma via multiple signal pathways. Sci Rep. 2016;6(July):1–9. doi:10.1038/srep37499

47. Lowery JW, de Caestecker MP. BMP signaling in vascular development and disease. Cytokine Growth Factor Rev. 2010;21(4):287–298. doi:10.1016/j.cytogfr.2010.06.001

48. Kamekura S, Kawasaki Y, Hoshi K, et al. Contribution of runt-related transcription factor 2 to the pathogenesis of osteoarthritis in mice after induction of knee joint instability. Arthritis Rheum. 2006;54(8):2462–2470. doi:10.1002/art.22041

49. Komori T, Yagi * H., Nomura S, Yamaguchi A. Targeted disruption of. Proc Natl Acad Sci U S A. 1998;95(15):8692–8697.

50. Meng X, Ziadlou R, Grad S, et al. Animal Models of Osteochondral Defect for Testing Biomaterials. Biochem Res Int. 2020;2020. doi:10.1155/2020/9659412

51. Sekiya I, Vuoristo JT, Larson BL, Prockop DJ. In vitro cartilage formation by human adult stem cells from bone marrow stroma defines the sequence of cellular and molecular events during chondrogenesis. Proc Natl Acad Sci U S A. 2002;99(7):4397–4402. doi:10.1073/pnas.052716199

52. Pelttari K, Winter A, Steck E, et al. Premature induction of hypertrophy during in vitro chondrogenesis of human mesenchymal stem cells correlates with calcification and vascular invasion after ectopic transplantation in SCID mice. Arthritis Rheum. 2006;54(10):3254–3266. doi:10.1002/art.22136

53. Mueller MB, Tuan RS. Functional characterization of hypertrophy in chondrogenesis of human mesenchymal stem cells. Arthritis Rheum. 2008;58(5):1377–1388. doi:10.1002/art.23370

54. Browe DC, Burdis R, Díaz-Payno PJ, et al. Promoting endogenous articular cartilage regeneration using extracellular matrix scaffolds. Mater Today Bio. 2022;16. doi:10.1016/j.mtbio.2022.100343

55. Steele JAM, Moore AC, St-Pierre JP, et al. In vitro and in vivo investigation of a zonal microstructured scaffold for osteochondral defect repair. Biomaterials. 2022;286. doi:10.1016/j.biomaterials.2022.121548

56. Park DY, Min BH, Park SR, et al. Engineered cartilage utilizing fetal cartilage-derived progenitor cells for cartilage repair. Sci Rep. 2020;10(1):1–13. doi:10.1038/s41598-020-62580-0

57. Zeng J, Huang L, Xiong H, et al. Injectable decellularized cartilage matrix hydrogel encapsulating urine-derived stem cells for immunomodulatory and cartilage defect regeneration. NPJ Regen Med. 2022;7(1):1–12. doi:10.1038/s41536-022-00269-w

58. Golchin A, Shams F, Karami F. Advancing Mesenchymal Stem Cell Therapy with CRISPR/Cas9 for Clinical Trial Studies. In: Cell Biology and Translational Medicine,. Vol 8.; 2019:89–100. doi:10.1007/5584_2019_459

59. Hazrati A, Malekpour K, Soudi S, Hashemi SM. CRISPR/Cas9-engineered mesenchymal stromal/stem cells and their extracellular vesicles: A new approach to overcoming cell therapy limitations. Biomedicine & Pharmacotherapy. 2022;156:113943. doi:10.1016/j.biopha.2022.113943

60. Chen Z, Klein T, Murray RZ, et al. Osteoimmunomodulation for the development of advanced bone biomaterials. Materials Today. 2016;19(6):304–321. doi:10.1016/j.mattod.2015.11.004

61. Xie Y, Hu C, Feng Y, et al. Osteoimmunomodulatory effects of biomaterial modification strategies on macrophage polarization and bone regeneration. Regen Biomater. 2020;7(3):233–245. doi:10.1093/rb/rbaa006

62. Aamodt JM, Grainger DW. Extracellular matrix-based biomaterial scaffolds and the host response. Biomaterials. 2016;86:68–82. doi:10.1016/j.biomaterials.2016.02.003

63. Ramakrishnaiah Siddappa, Ruud Licht, Clemens van Blitterswijk J de B. Donor variation and loss of multipotency during in vitro expansion of human mesenchymal stem cells for bone tissue engineering. Journal of Orthopaedic Research. 2007;25(8).

64. LaFleur MW, Nguyen TH, Coxe MA, et al. A CRISPR-Cas9 delivery system for in vivo screening of genes in the immune system. Nat Commun. 2019;10(1):1–10. doi:10.1038/s41467-019-09656-2

65. Bock C, Datlinger P, Chardon F, et al. High-content CRISPR screening. Nature Reviews Methods Primers. 2022;2(1). doi:10.1038/s43586-021-00093-4

66. Maehara H, Sotome S, Yoshii T, et al. Repair of large osteochondral defects in rabbits using porous hydroxyapatite/collagen (HAp/Col) and fibroblast growth factor-2 (FGF-2). Journal of Orthopaedic Research. 2010;28(5):677–686. doi:10.1002/jor.21032

